# Population level gene expression can repeatedly link genes to functions in maize

**DOI:** 10.1101/2023.10.31.565032

**Authors:** J. Vladimir Torres-Rodríguez, Delin Li, Jonathan Turkus, Linsey Newton, Jensina Davis, Lina Lopez-Corona, Waqar Ali, Guangchao Sun, Ravi V. Mural, Marcin W. Grzybowski, Addie M. Thompson, James C. Schnable

## Abstract

Transcriptome-Wide Association Studies (TWAS) can provide single gene resolution for candidate genes in plants, complementing Genome-Wide Association Studies (GWAS) but efforts in plants have been met with, at best, mixed success. We generated expression data from 693 maize genotypes, measured in a common field experiment, sampled over a two-hour period to minimize diurnal and environmental effects, using full-length RNA-seq to maximize the accurate estimation of transcript abundance. TWAS could identify roughly ten times as many genes likely to play a role in flowering time regulation as GWAS conducted data from the same experiment. TWAS using mature leaf tissue identified known true positive flowering time genes known to act in the shoot apical meristem, and trait data from new environments enabled the identification of additional flowering time genes without the need for new expression data. eQTL analysis of TWAS-tagged genes identified at least one additional known maize flowering time gene through *trans*-eQTL interactions. Collectively these results suggest the gene expression resource described here can link genes to functions across different plant phenotypes expressed in a range of tissues and scored in different experiments.

## Introduction

Information from homologous genes can predict the molecular functions of the proteins encoded by genes with reasonably high accuracy (e.g. which genes are transcription factors and which are transmembrane transporters). However, there can be significant variability in determining the specific biological processes in which homologous proteins participate and contribute. Even in the most widely studied plant genetic models – maize, rice, and Arabidopsis – only a modest proportion of annotated gene models (1-10%) have been directly linked to their roles in determining plant phenotypes (Schnable and Freeling 2011; Lloyd and Meinke 2012). This lack of direct functional information on the role individual genes play in determining plant phenotypes is even more striking for agricultural crops and wild species which have historically not served as genetic models (Rhee and Mutwil 2014; Boyles *et al*. 2019). The advent of gene editing technology has accelerated this process, but the functional characterization of a single gene’s role in determining plant phenotype continues to require substantial investments of both time and resources. As a result, it is likely that the vast majority of annotated gene models will continue to lack this gold standard functional information for the foreseeable future.

Genome-wide association studies (GWAS) that link genetic markers to variation in plant phenotypes have been widely adopted in plant species (as reviewed by (Tibbs Cortes *et al*. 2021)). These studies can act as a partial substitute for characterizing the function of specific genes through loss of function alleles or as a method for prioritizing candidate genes for subsequent functional characterization. Such characterization is typically time-consuming and often yields results that fall short of expectations. GWAS approaches also have several key limitations. The first is that genes vital to a phenotype of interest will not be identified in a GWAS if functional variation for the gene of interest is not present in the studied population, or present only at low frequencies that reduce statistical power to discover variants. The second key limitation of GWAS is that the signals it identifies tag regions of the genome linked to variation in a phenotype, but these regions can include multiple annotated gene models. As a result, it is frequently not possible to conclude which specific gene is responsible for a given GWAS signal without additional time and resource-intensive follow-up experiments.

Transcriptome-wide association studies (TWAS) test for significant associations between the expression of individual genes and variation in plant phenotypes. TWAS partially addresses both key limitations of GWAS described above. TWAS identifies complementary rather than redundant sets of genes to those identified via GWAS for the same phenotypes in the same populations. The expression level of an individual gene can integrate the signals from multiple upstream regulatory variants, each too small or too rare to be linked directly to variation in the phenotype of interest (Li *et al*. 2023). TWAS based on direct measurements of gene expression typically identifies specific candidate genes rather than intervals containing multiple genes, even in species or populations with slow decay of linkage disequilibrium across the genome (Li *et al*. 2023). However, the terminology TWAS has also been applied to methodologies that use genetic marker data to impute gene expression and seek to link that imputed gene expression data to phenotype. In these cases, the advantage of single gene resolution that TWAS provides is lost (Wainberg *et al*. 2019; Mai *et al*. 2023).

Several challenges have limited the widespread application of transcriptome-wide association in plants. A large proportion of plant transcripts exhibit diurnal cycling including >90% of transcripts in Arabidopsis (Michael *et al*. 2008), 60% of transcripts in rice and poplar (Filichkin *et al*. 2011) and 30-50% of transcripts in maize, sorghum, and foxtail millet (Lai *et al*. 2020). Given the need to flash freeze tissue to avoid wound-induced changes to gene expression, it can be difficult to sample sufficiently large populations in short enough periods of time to avoid the confounding effects of diurnal changes in gene expression. The size of populations required for successful TWAS analyses also presents financial barriers to the use of this method, due to the high cost of RNA sequencing relative to many low-cost DNA genotyping technologies. To partially mitigate this issue, 3 ’ tail RNA-seq can be used and it has been shown to have similar levels of repeatability to whole transcript RNA sequencing but has some disadvantages, such as the reduced ability to detect differentially expressed genes, especially for long transcripts and the loss of information in the 5 ’ end of the transcript (Ma *et al*. 2019), plus don’t provide splicing information in case isoform level analysis or genome-wide association study is needed.

The combined impact of the above factors is that TWAS studies in plants are often limited in size, reducing statistical power. They employ data collected from different genotypes at different times, thereby increasing non-genetic variation in gene expression, or employing lower-cost techniques that profile the expression of fewer genes at lower resolution. As a result, in many cases, TWAS has not identified any genes above rigorous false discovery thresholds and instead must use approaches such as considering the top 1% of most significantly associated genes (Kremling *et al*. 2019). This approach provides significant biological insight but is likely to include some proportion of false positive associations alongside true biological signals, again adding complexity to pinpointing the actual causal genes.

A key logistical advantage of GWAS is that, while generating high density resequencing data can be as expensive or more expensive than profiling gene expression across the entire genome, once a specific association population has been genotyped, the same genetic marker data can be employed multiple times to map different traits of interest allowing the high cost of data to be amortized across many research projects. In contrast, gene expression patterns change across tissues and developmental stages, as well as in response to changes in the environment. This raises concerns about how much, if any, potential exists to reuse transcriptome-wide expression datasets to identify genes linked to variation in different traits in different environments. Multiple studies have demonstrated that gene expression from non-target tissues and non-target environments can identify true positive causal genes (Hirsch *et al*. 2014; Li *et al*. 2021, 2023). However, the reuse of transcript abundance data across multiple studies conducted using the same population remains less widely adopted than the reuse of genetic marker data across studies.

Here we sought to generate a reusable set of transcript abundance measurements for the expanded Wisconsin Diversity panel, a large panel of temperate adapted maize lines (Mazaheri *et al*. 2019). We evaluated the power and accuracy of this dataset to identify genes of interest using flowering time data collected from two different environments. TWAS identified approximately 10 times as many positive hits as GWAS conducted using the same trait datasets. The genes identified via TWAS included many known true positive flowering time genes missed by GWAS, genes linked to flowering time regulation in rice or Arabidopsis but not in maize, and a modest number of genes previously unlinked to flowering time but with plausible functional mechanisms connecting them to this trait. Notably, the genes identified via TWAS conducted using expression in mature leaf tissue include genes shown to act primarily or exclusively in the shoot apical meristem. The expression quantitative trait loci (eQTL) analysis conducted using TWAS flowering time hits further extended gene discovery, including at least one additional known true positive maize flowering time gene which was not directly tagged by TWAS but identified as a *trans*-eQTL regulator. Overall, these results demonstrate both the potential of TWAS for assigning putative functions to genes and the potential to reuse gene expression datasets to analyze traits collected in multiple environments.

## Results

Maize is a monoecious species with separate specialized male and female flowers. As a result, two separate flowering time phenotypes can be scored for maize: one based on the time when anthers emerge from the tassel – the specialized male inflorescence – and a second based on the time when silks emerge from the ear – the specialized female inflorescence. Male and female flowering times were scored across replicated field experiments grown in Michigan and Nebraska in 2020. Both male (hereafter referred to as anthesis) and female (hereafter referred to as silking) flowering occurred fewer days after planting in Michigan than in Nebraska (Supplemental dataset S1 & S2). The mean number of days to anthesis in Michigan was 64, while in Nebraska it was 72. The mean number of days to silking in Michigan was 67, while in Nebraska it was 75 (Supplemental Figure S1). The within-environment repeatability was high for the four trait datasets, although modestly higher for anthesis (0.90 in Michigan, 0.87 in Nebraska) than for silking (0.87 in Michigan, 0.84 in Nebraska).

Mature leaf tissue was sampled from 750 plants in the Nebraska field experiment over a two-hour period (Figure 1A). A median of 83% of the RNA sequence reads generated via RNA-seq of RNA extracted from these tissue samples could be uniquely assigned to the primary transcript of one of the 39,756 annotated protein-coding gene models from the B73_RefGen_V5 reference genome (Hufford *et al*. 2021). A total of 699 unique genotypes were represented among the 750 plants, with 51 genotypes represented twice. Two samples were excluded as outliers based on the results of a PCA conducted using transcripts per million (TPM) based estimates of gene expression values (Supplemental Figure S2). Four additional samples were excluded from downstream analysis because they were either not included in a recent resequencing-based genetic marker dataset (DK84QAB1, HP72-11, PHT69), or were expected to be largely isogenic with another sample (B73Htrhm was excluded while B73 was retained) (Grzybowski *et al*. 2023). Maize genes with an average expression >4 TPM exhibited an average repeatability of *∼*0.6 estimated expression across genetic replicates. Average repeatability of gene expression declined for genes with average expression levels <4 TPM, plateauing at *∼*0.4 with an average expression of 0.1 TPM (Supplemental Figure S3). Two main clusters of genes were observed: one of genes with extremely low repeatability (0.0 - 0.05) centered on an average expression of approximately 0.01 TPM, and a second of higher repeatability genes (0.6 - 0.8) with average expression values between 1 and 100 TPM (Supplemental Figure S3). The set of 24,585 gene models with expression *≥* 0.1 TPM in at least 347 of the remaining 693 genotypes were retained for subsequent analyses.

**Figure 1.**
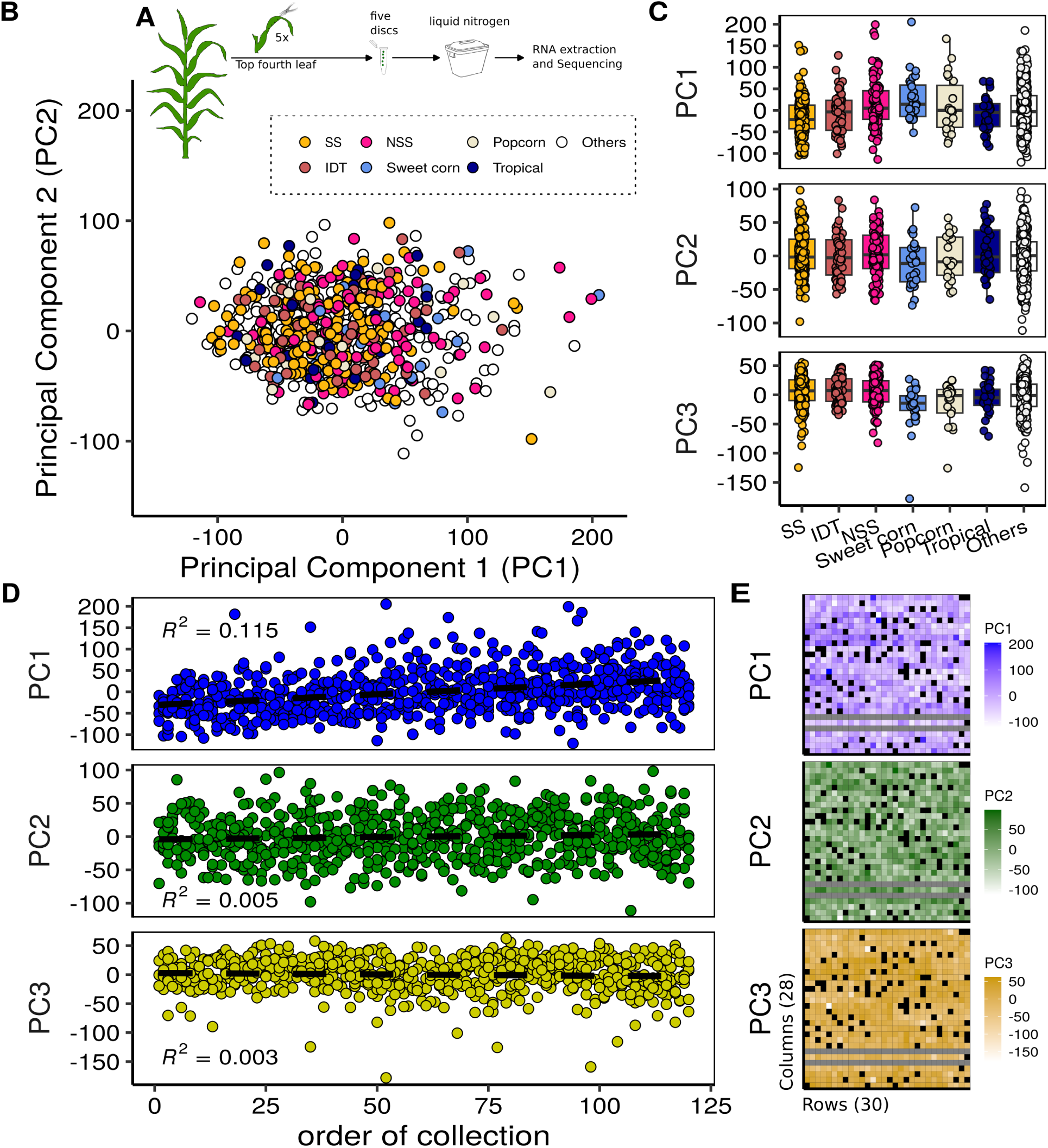
Sources of potential confounding variation in expression data. **A** Methodology employed to collect tissue samples for gene expression profiling **B** Principal component (PC) scores for 693 RNA-seq libraries representing unique maize genotypes, classified using the subpopulation assignments reported in Grzybowski *et al*. (2023). SS = the stiff stalk heterotic group. NSS = non-stiff stalk heterotic group. IDT = iodent heterotic group. PC1 and PC2 explain 10.8% and 4.8% of total variation in gene expression respectively. **C** Distribution of scores for the first three PCs among maize genotypes assigned to each population. **D** Relationships between the first three PCs and order of sample collection. Samples were collected by several researchers in parallel so multiple samples share the same sample order value. Dashed black line indicates the best fit linear regression. **E** Relationships between the first three PCs and the spatial distribution of sampled plants across the field. Grey boxes indicate plots not sampled. Black boxes indicate either check plots (skipped), or samples excluded at the quality control stage.

The non-repeatable component of variation in gene expression between replicated samples of the same maize genotypes in the same field can result from a number of factors, including diurnal cycling of gene expression and micro-environmental variation across the field. The first three principal components (PCs) of variation in gene expression among the 24,585 genes in 693 maize genotypes that passed filtering criteria explained 10.8 %, 4.78 %, and 3.42 % of the total variation in the dataset (Figure 1B). While differences in the distribution of PC values existed between some maize sub-populations, distributions of PC scores were largely overlapping (Figure 1B,C). Order of sample collection, a proxy for time of collection, was recorded for all samples, which allowed us to measure its impact on gene expression. Order of collection was most correlated with PC5 (percent of variance in gene expression explained=2.4%, R^2^ with collection order=0.18), PC1 (percent of variance in gene expression explained=10.8%, R^2^ with collection order=0.11), and PC7 (percent of variance in gene expression explained=1.9%, R^2^ with collection order=0.06). The R^2^ of all other PCs with the order of collection was <0.02 (Figure 1D and Supplemental dataset S3). As a positive control, we examined the expression of four core maize circadian clock genes (Lai *et al*. 2020) representing two clock components expected to be decreasing or increasing in expression at the time of collection: *lhl1* (R^2^ = 0.25) & *lhl2* (R^2^ = 0.39) and *gi1* (R^2^ = 0.25) & *gi2* (R^2^ = 0.20) (Supplemental Figure S4). The correlation of *lhl2* with order of collection was the second highest of any gene in the dataset. Overall, few genes (*∼*4.2%) used in downstream analysis exhibited a correlation R^2^ higher than 0.1 with order of collection (Supplemental Figure S5). None of the top ten PCs were correlated with row or column positions above an R^2^=0.03 (Supplemental dataset S3), furthermore, no other obvious non-linear associations were observed between PCs and field layout (Figure 1E).

Transcriptome-wide association studies (TWAS) conducted using the filtered gene expression data collected from Nebraska along with flowering time traits (days to anthesis and days to silking) scored in both Nebraska and Michigan environments identified 21 unique gene-trait associations at a false discovery rate threshold (FDR) of 5%. Between 15 and 18 genes were identified per individual trait. A set of 12 genes were consistently identified across all four analyses (Figure 2 and Table 1). Three of the 21 genes identified by TWAS have been previously shown to alter flowering time in maize: *zmm4* (Figure 3A), a MADS-box transcription factor that functions downstream of *zcn8* in the shoot apical meristem (Danilevskaya *et al*. 2008); *zcn8* (Figure 3B), thought to act as the mobile florigen from leaves to the shoot apical meristem (Meng *et al*. 2011); and *mads1* (Figure 5B), an ortholog of the floral integrator *SUPPRESSOR OF OVEREXPRESSION OF CONSTANS1* (*soc1*) (Alter *et al*. 2016). In an additional ten cases, the rice or Arabidopsis orthologs of the maize genes identified via TWAS had been linked to variation in flowering time (Table 1). A conventional genome-wide association analysis (GWAS) conducted with the same trait datasets and the same population identified only two consistent/strong signals: one localized near *zcn8*, and a second near *mads69*, a major flowering time locus in maize validated via loss of function alleles (Liang *et al*. 2019) (Supplemental Figure S6).

**Figure 2.**
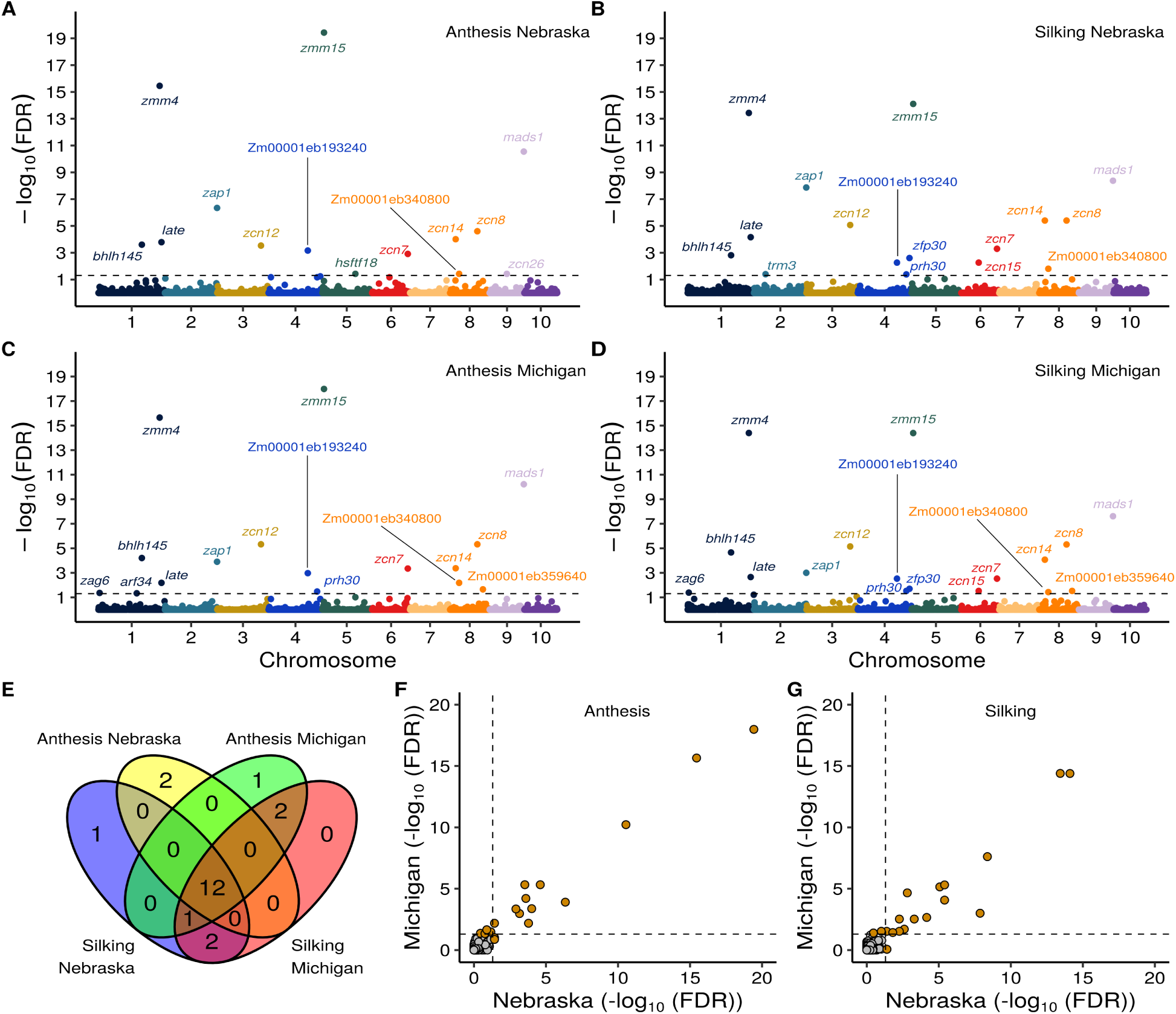
Genes associated with variation in anthesis or silking time in Nebraska and Michigan via transcriptome-wide association. **A** Results of a transcriptome-wide association study conducted using transcript abundance data and spatially corrected days to anthesis measured in Nebraska in 2020. Horizontal dashed line indicates a 0.05 False Discovery Rate (FDR) cutoff as determined by the Benjamini–Hochberg method. **B** Results of a transcriptome-wide association study conducted using transcript abundance data and spatially corrected days to silking measured in Nebraska in 2020. Plotted as described in panel A. **C** Results of a transcriptome-wide association study conducted using transcript abundance data and spatially corrected days to anthesis in Michigan in 2020. Plotted as described in panel A. **D** Results of a transcriptome-wide association study conducted using transcript abundance data and spatially corrected days to silking in Michigan in 2020. Plotted as described in panel A. **E** Numbers of shared and uniquely identified genes in the four TWAS results presented in panels A-D. **F** Relationship between FDRs assigned to the same genes in TWAS conducted using anthesis measurements in Nebraska and Michigan. Dashed lines indicate 0.05 FDR cutoffs. **G** Relationship between FDRs assigned to the same genes in TWAS conducted using silking measurements in Nebraska and Michigan. Dashed lines indicate 0.05 FDR cutoffs.

**Figure 3.**
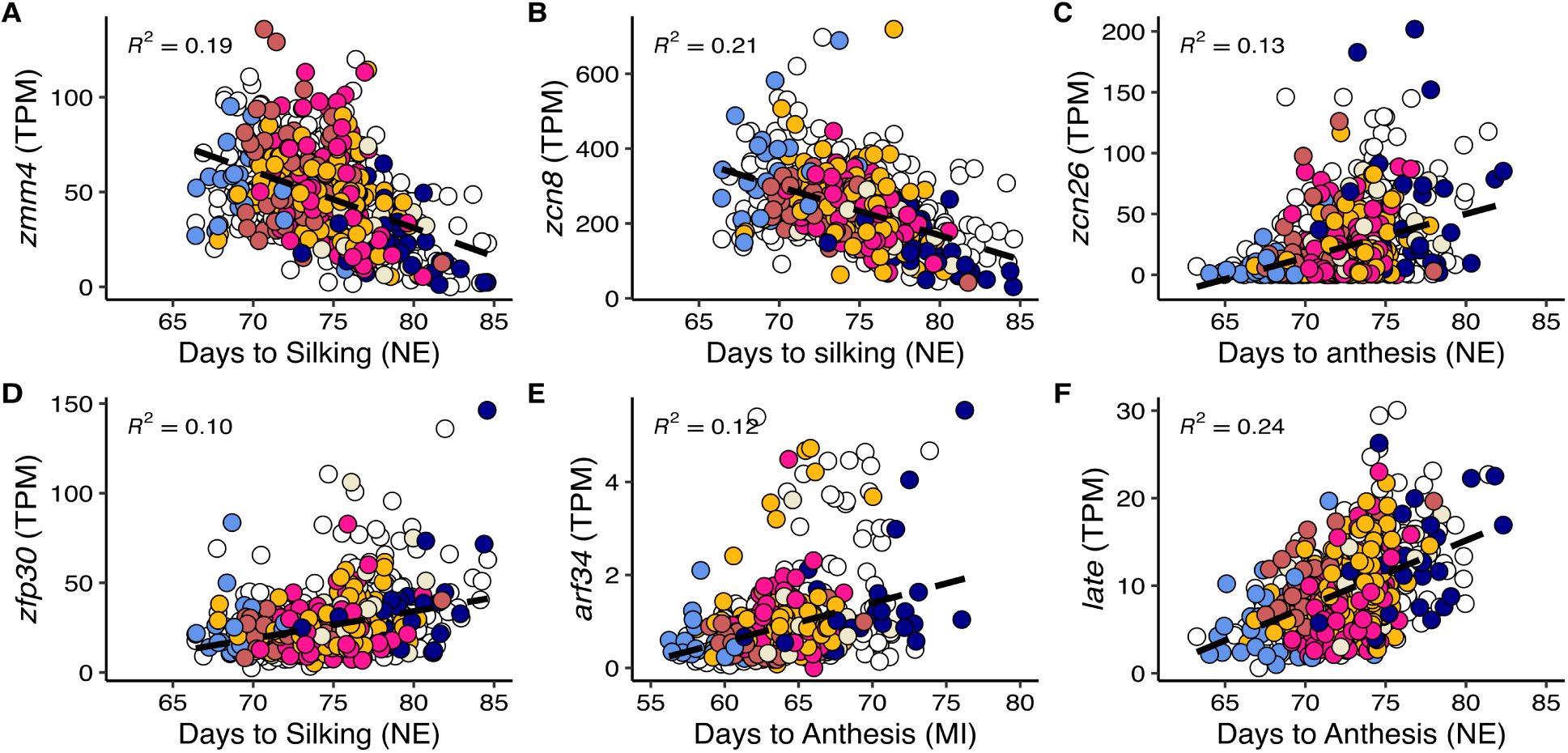
Relationships between gene expression and flowering time for a subset of significant genes identified via TWAS. **A** Relationship between the expression of *zmm4* in mature leaf tissue of different maize genotypes and days to silking in Nebraska for the same genotypes. Colors indicate subpopulation assignments from Figure 1B. Black dashed line indicates linear best fit. **B** Relationship between the expression of *zcn8* in different maize genotypes and and days to silking in Nebraska. **C** Relationship between the expression of *zcn26* in different maize genotypes and and days to anthesis in Nebraska. **D** Relationship between the expression of *zfp30* in different maize genotypes and and days to silking in Nebraska. **E** Relationship between the expression of *arf34* in different maize genotypes and and days to anthesis in Michigan. **F** Relationship between the expression of *late* in different maize genotypes and and days to anthesis in Nebraska.

**Figure 4.**
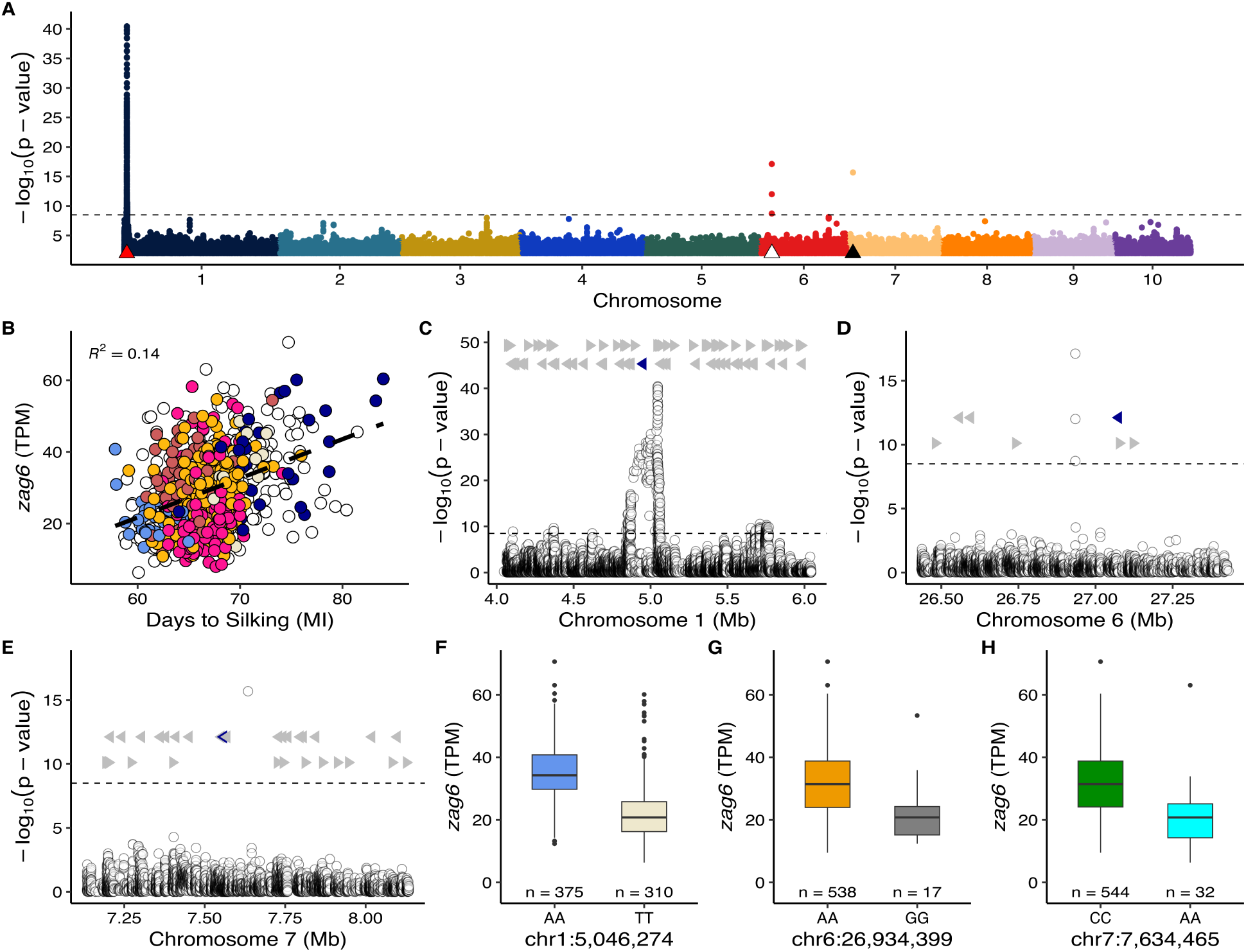
Details of several eQTLs associated with the expression of *zag6*. **A** Genome-wide association identifying genetic markers linked to variation in the expression of *zag6*. The red triangle at the bottom of the dots in chromosome 1 indicates the position of *zag6*. White and black triangles at the bottom of the dots in chromosomes 6 and 7 indicate the postions of *lbl1* and *hen1*, respectively. Black horizontal dashed line indicates a Bonferroni corrected 0.05 significance threshold. **B** Relationship between *zag6* expression and days to silking scored from Michigan 2020. Dots are colored based on sub-groups referred to in Figure 1. Linear dashed lines indicate the linear regression fit to this data. **C** Zoomed in view of the region on chromosome 1 containing the *cis*-eQTL for *zag6*. Black horizontal dashed line indicates Bonferroni corrected 0.05 significance threshold. Triangles indicate the positions of annotated genes in the region. The blue triangle indicates the annotated position of *zag6* specifically. **D** Zoomed in view of the region on chromosome 6 containing a *trans*-eQTL for *zag6*.Blue triangle indicates the position of *lbl1*. **E** Zoomed in view of the region on chromosome 7 containing a *trans*-eQTL for *zag6*.Blue triangle indicates the position of *hen1*. **F** Effect of the most significantly associated genetic marker in the *cis*-eQTL on the expression of *zag6*. **G** Effect of the most significantly associated genetic marker in the *trans*-eQTL on on chromosome 6 the expression of *zag6*. **H** Effect of the most significantly associated genetic marker in the *trans*-eQTL on on chromosome 7 the expression of *zag6*.

**Figure 5.**
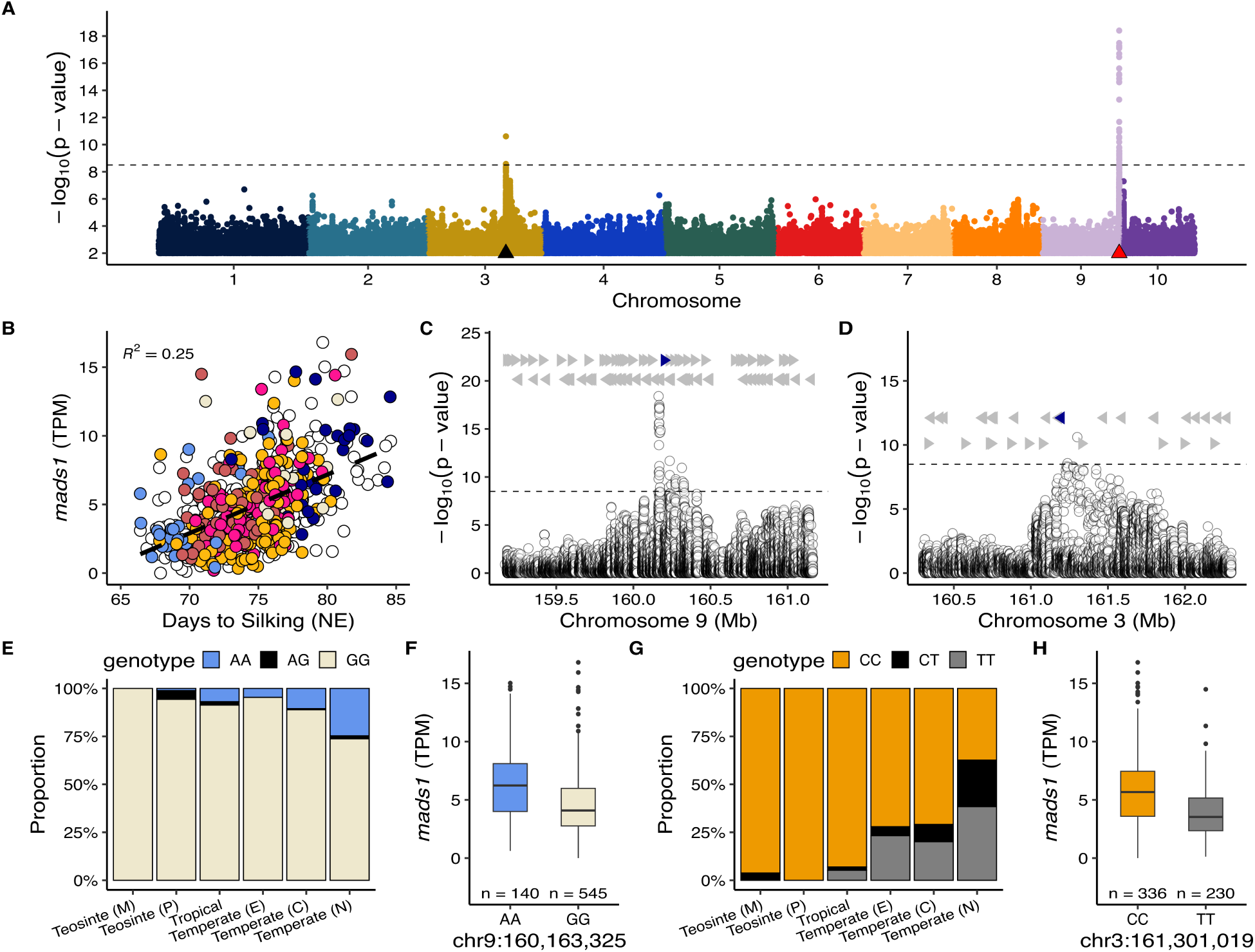
Localization and changes in allele frequencies of eQTLs associated expression of *mads1*. **A** Genetic markers associated with variation in the expression of *mads1*. Red triangle marks the position of *mads1*. Black triangle marks the position of *mads69*. Black horizontal dashed line indicates a 0.05 threshold after Bonferroni correction. **B** Relationship of *mads1* expression to days to silking scored in Nebraska. Dots are colored based on sub-groups referred to in Figure 1. The dashed line indicates the calculated linear regression. **C** Zoomed in view of the peak located on chromosome 9. Triangles indicate the position of annotated genes in the region. Blue triangle indicates the position of *mads1*. **D** Zoomed in view of the peak located on chromosome 3. Blue triangle indicates the position of *mads69*. **E** Variation in the frequencies of different alleles of the most significantly associated genetic marker within the peak on chromosome 9 across different wild and domesticated maize populations (Supplemental dataset S4). **F** Differences in the expression of *mads1* between maize genotypes carrying different alleles the genetic marker shown in panel E. **G** Variation in the frequencies of different alleles of the most significantly associated genetic marker within the peak on chromosome 3 across different wild and domesticated maize populations. **H** Differences in the expression of *mads1* between maize genotypes carrying different alleles the genetic marker shown in panel E.

**Table 1.** Associated genes via Transcriptome-Wide association study from different environments using a single gene expression dataset.

| Gene ID | Symbol | *cis-eQTL* | *trans-eQTL* | Avg. Exp. | Environment | Protein Function |
| --- | --- | --- | --- | --- | --- | --- |
| Zm00001eb214750 | zmm15 |  | yes | 49.09 | A.NE, S.NE, A.MI, S.MI | MADS box TF/AGL79-like |
| Zm00001eb057540 | zmm4 |  |  | 46.81 | A.NE, S.NE, A.MI, S.MI | MADS box TF/AGL79-like |
| Zm00001eb403750 | mads1 | yes | yes | 4.93 | A.NE, S.NE, A.MI, S.MI | MADS box TF/SOC1-like |
| Zm00001eb118120 | zap1 |  | yes | 2.96 | A.NE, S.NE, A.MI, S.MI | MADS box TF/AGL79-like |
| Zm00001eb338650 | zcn14 |  |  | 1.24 | A.NE, S.NE, A.MI, S.MI | PEBP/FT-like |
| Zm00001eb353250 | zcn8 |  |  | 236.59 | A.NE, S.NE, A.MI, S.MI | PEBP/FT-like |
| Zm00001eb153190 | zcn12 |  | yes | 176 | A.NE, S.NE, A.MI, S.MI | PEBP/FT-like |
| Zm00001eb059970 | late | yes | yes | 9.3 | A.NE, S.NE, A.MI, S.MI | Zinc finger TF/C2H2 |
| Zm00001eb293080 | zcn7 |  |  | 228.84 | A.NE, S.NE, A.MI, S.MI | PEBP/FT-like |
| Zm00001eb037440 | bhlh145 | yes | yes | 0.48 | A.NE, S.NE, A.MI, S.MI | bHLH TF |
| Zm00001eb205550 | zfp30 |  |  | 26.13 | S.NE, S.MI | RNA binding zinc-finger |
| Zm00001eb193240 | – |  |  | 13.99 | A.NE, S.NE, A.MI, S.MI | Zinc finger TF/C2H2 |
| Zm00001eb271180 | zcn15 |  |  | 1.66 | S.NE, S.MI | PEBP/FT-like |
| Zm00001eb340800 | – | yes |  | 2.04 | A.NE, S.NE, A.MI, S.MI | Unknown function |
| Zm00001eb082790 | trm3 | yes | yes | 8.07 | S.NE | Thioredoxin |
| Zm00001eb203040 | prh30 |  |  | 9.1 | S.NE, A.MI, S.MI | Protein phosphatase |
| Zm00001eb001670 | zag6 | yes | yes | 29.34 | A.MI, S.MI | MADS box TF/SOC1-like |
| Zm00001eb031700 | arf34 | yes | yes | 0.89 | A.MI | Auxin response factor |
| Zm00001eb239380 | hsftf18 | yes |  | 3.44 | A.NE | Heat Shock TF |
| Zm00001eb359640 | – | yes |  | 44.71 | A.MI, S.MI | Unknown function |
| Zm00001eb384770 | zcn26 | yes | yes | 22.48 | A.NE | PEBP/FT-like |

PEBP = Phosphatidyl ethanolamine-binding protein PEBP, A.NE = Anthesis in Nebraska, S.NE = Silking in Nebraska, A.MI = Anthesis in Michigan, S.MI = Silking in Michigan. Avg. Exp = Average expression reported as Transcripts per million

Multiple members of several gene families were present among the set of 21 flowering time TWAS hits. These included six members of the *Zea mays CENTRORADIALIS* (*ZCN*) family (*zcn7*, *zcn8*, *zcn12*, *zcn14*, *zcn15* and *zcn26*) and five MADS-box transcription factors including three members of the AGL-79-like subgroup (*zmm4*, *zmm15*, *zap1*) and two members of the SOC1-like subgroup (*mads1* and *zag6*). Several gene families were represented by multiple family members. In three cases, these represent homeologous gene copies from the maize whole genome duplication: *mads1* and *zag6*, *zmm4* and *zmm15*, and *zcn7* and *zcn8*. The gene pairs *zmm4* and *zmm15* & *zcn7* and *zcn8* were the two pairs of genes among our hits whose expression was most correlated, while the expression of *mads1* and *zag6* was more diverged (Supplemental Figure S7). Greater expression of five of the six *zcn* genes identified was associated with earlier flowering time, while greater expression of *zcn26* was associated with later flowering time (R^2^ = 0.13, Figure 3B,C). Notably, *zcn26* is also one of the six *zcn* genes identified where transgenic expression failed to rescue the delayed flowering time phenotype of the *ft* mutant in Arabidopsis (Stephenson *et al*. 2019) while retaining the capacity to interact with the floral transition promoter DLF1 protein (Meng *et al*. 2011), suggesting this gene may have the capacity to act as competitive inhibitor of the flowering activating complex.

There are at least two potential explanations for why TWAS can identify true positive gene-trait associations not identified in GWAS conducted with the same populations and phenotype data. One is that the expression of individual genes can reflect and integrate the impact of multiple *trans* regulatory variants which may not individually have sufficient effect sizes or minor allele frequencies to be detected in GWAS for the target trait. A second is that the expression of a gene can capture the effects of three or more functionally variable *cis* regulatory haplotypes (defined by two or more SNPs) with different effects on gene expression and the target phenotype, information which would be missed when testing for association with individual biallelic genetic markers, which can fail to capture variance from large number of variable haplotypes.

A number of both *cis* and *trans*-eQTL were identifiable among the genes initially linked to flowering time via TWAS. In three cases, the expression genes identified via TWAS for flowering time were also associated with one or more *cis*-eQTL but no *trans*-eQTL. In another three cases, the expression of genes identified via TWAS for flowering time was associated with one or more *trans*-eQTL but no *cis*-eQTL. In seven cases the expression of genes identified via TWAS was associated with both an eQTL acting in *cis* and one or more eQTL acting in *trans* (Table 1).

The set of genes where both *cis* and *trans*-eQTL were identified included AGAMOUS-LIKE6 *zag6*, a homolog of the flowering time integrator *soc1* in Arabidopsis. Higher expression of *zag6* was associated with greater numbers of days to anthesis and days to silking in Michigan but not in Nebraska (Figure 2 & 4B). The single most significant SNP within a *cis*-eQTL associated with *zag6* was located 98.1 Kb upstream of the gene’s transcription start site (Figure 4C). Two additional signals associated with the expression of *zag6* were located in *trans*, one on chromosome 6 (chr6:26,934,399) and the other in chromosome 7 (chr7:7,634,465) (Figure 4A). Both signals are associated with genes involved in small RNAs. The closest gene to the chromosome 6 signal is *leafbladeless1* (*lbl1*, 134 kb), a gene involved in the trans-acting short-interfering RNA (ta-siRNA) biogenesis pathway (Dotto *et al*. 2014) while the second closest gene to the chromosome 7 signal is *hen1* (81 kb), a small RNA methyltransferase involved in processing small RNAs (Figure 4C-H) (Park *et al*. 2002; Xie *et al*. 2004; Yu *et al*. 2005).

The gene *mads1* has been reported to function as a floral activator in maize (Alter *et al*. 2016). However, while *mads1* was identified as significantly linked to variation in both male and female flowering time in both Nebraska and Michigan (Figure 2), the expression of *mads1* was significantly negatively correlated with flowering (Figure 5B & Supplemental Figure S8). A genome-wide association study conducted for genetic markers linked to variation in the expression of *mads1* identified two significant signals (Figure 5A). The first appears to be a *cis* regulatory variant, with the most significant SNP of the peak located 47 kilobases upstream from *mads1* on chromosome 9 (Figure 5C). The second peak associated with the expression of *mads1* is located on chromosome 3, in the distal promoter region of *mads69* (Figure 5D), the one example of a well characterized flowering time gene in maize which was identified via GWAS in this population, but not via TWAS (Liang *et al*. 2019). The expression of two additional genes identified via TWAS for flowering time (*zmm15* and *zap1*) were also significantly associated with *trans*-eQTL in the vicinity (89.2 and 51.6 kilobases away respectively) of *mads69* (Supplemental Figure S9). The peaks of these three trans-eQTL define a total interval of 63,397 base pairs and are all in reasonably high linkage disequilibrium with each other (R^2^=0.77-0.92).

The SNP most significantly linked to variation in the expression of *mads1* within the promoter of *mads69* showed significant shifts in allele frequency between wild, domesticated tropical, and domesticated temperate maize populations (Figure 5G). The allele associated with reduced expression of *mads1* 5H) was extremely rare in wild teosinte populations and never observed in a homozygous state among sampled representatives of those populations. The frequency of this allele increases moderately in tropical domesticated maize, and further in temperate domesticated maize populations on three continents (Figure 5G). These shifts are consistent with the reported signature of selection previously described in the region of *mads69* (Liang *et al*. 2019). However, a compensatory pattern of allele frequency change was observed at the *mads1 cis*-regulatory variant. Here the allele associated with increased expression of *mads1* 5C) is the allele which is observed only at extremely low frequencies among wild teosinte samples, but is found at increasing frequency in domesticated populations, particularly those from temperate regions 5E,F)

## Discussion

Transcriptome-Wide association studies have been explored a number of times in plants with mixed results. In some cases only one or several genes are significant above false discovery thresholds (Hirsch *et al*. 2014; Lin *et al*. 2017). In many others, the top 0.5-1% of genes are evaluated rather than applying multiple testing correction, an approach that can enrich for true positives but likely with significant proportions of false positives included as well (Kremling *et al*. 2019; Wu *et al*. 2022). This choice is likely made because, in these cases, there were no individual genes associated with the target phenotype at significance thresholds that rigorously control false discovery rates. Here we report transcriptome-wide association studies identifying a total of 21 genes surpassing stringent false discovery rate thresholds. Many of these genes can be validated based on existing literature reports, relative to only two genes identified via genome-wide association using the same dataset. However, applying standard Bonferroni correction to TWAS only modestly reduced the number of genes passing statistical significance thresholds (11 significantly associated genes via Bonferroni vs 14 significant genes via Benjamini-Hochberg control, Figure 2A & Supplemental Figure S10). Other factors which may explain the greater power we observe to discover phenotype associated genes in this study relative to previous TWAS may include the greater size of the population analyzed, the short length of time allowed for all sample collection, and the use of full length RNA-seq rather than three prime tail seq. Even within our two-hour sampling window, significant changes in the expression of diurnally cycling genes were observed (Supplemental Figure S4) and many genes exhibited modest correlations with sampling order (Figure 1D). Conducting sample collection over longer time frames or across multiple days would almost certainly exacerbate these sources of variation in measured gene expression, creating additional noise when seeking to link gene expression to variation in plant traits. Three prime tail seq has been preferred to conventional RNA-seq for TWAS applications given its lower cost per sample. However, it may be that, by targeting a single region of the 3 prime UTR, this method increases the probability that sequence polymorphisms between individuals reduce alignment rates to specific genes in specific individuals, creating variation in measured gene expression that reflects sequence differences rather than differences in the relative abundance of mRNA transcripts. The necessity of minimizing per sample costs is greater for datasets which can only be used once than for datasets which are reusable. We sought to evaluate whether the transcript abundance dataset we collected would only be usable for trait data collected in the same environment or if it could be reused for new trait datasets collected in new environments. The combination of Nebraska transcript data with Michigan flowering time data produced good results (Figure 2, Table 1). These included not only the re-identification of many of the same genes identified using trait data collected from the same field where transcript abundance was measured but also additional significantly associated genes, including one gene previously linked to a role in maize floral development (Figure 2, Table 1) suggesting at least some potential to discover useful new gene-trait associations by collecting new trait datasets in new environments without the need to generate new transcript datasets. However, another challenge is that different traits will be expressed in different tissues. While at least two previous studies suggest that known causal genes can be identified using transcripts from tissues other than those in which the trait is expressed, it is still quite notable that two of the genes whose expression in mature leaf tissue was most closely associated with flowering time – *zmm4* and *zmm15* – are genes known to act in the meristem and not previously thought to be expressed in mature leaf tissue (Danilevskaya *et al*. 2008). In addition to *zmm4* and *zmm15*, numerous other genes identified in this analysis have been previously linked to flowering time variation in maize (Table 1). While many genes were linked to both variation in days to anthesis and days to silking, two genes, *zfp30* (Figure 3D) and *zcn15*, were associated with days to silking in both environments but not with days to anthesis in either (Table 1). Greater expression of *zcn15*, which is syntenic with the the rice FT-like genes Hd3a and Hd3b (Tsuji *et al*. 2008), is associated with earlier silking. Two genes were also consistently associated with variation in both male and female flowering in Michigan, but not in Nebraska. These included *zag6*, a gene previously identified using a TWAS, there referred to as *zagl1*, conducted using measurements of flowering time recorded near Madison, Wisconsin (where termed *zagl1*) (Hirsch *et al*. 2014), a location at a very similar latitude to East Lansing, Michigan (43.1 N vs 42.7 N) but not to Lincoln, NE. One major flowering time gene which was notable by its absence from the TWAS results was *mads69*. The role of this gene in flowering time has been validated via loss of function studies (Liang *et al*. 2019), it has been detected in multiple genome-wide associations conducted using the same association population (Mazaheri *et al*. 2019; Grzybowski *et al*. 2023), and it has previously been linked to flowering time via TWAS conducted using gene expression data from early stage seedlings and other tissues (Hirsch *et al*. 2014; Lin *et al*. 2017; Li *et al*. 2021). One potential explanation was that *mads69* was not expressed in our target tissue, mature leaves. However, *mads69* exhibited a median expression level of 15 TPM, substantially higher than a number of other true positive genes identified via TWAS. Its expression in our dataset was simply not correlated with flowering time (Supplemental Figure S11). However, even in this case of a known true gene-trait association where it is clear our sampling occurred at the wrong time point and/or targeted the wrong tissue, this population scale transcript abundance data still recovered *mads69* as an eQTL hotspot associated with variation in the expression of three genes identified via TWAS which are presumably downstream of *mads69*: *mads1*, *zap1*, and *zmm15*. An analysis which recovers only genes already know to play roles in the trait of interest may be statistically sound, but ultimately does not contribute a great deal of additional knowledge about gene function. The analyses described above also identified genes not previously linked to flowering time in maize which may play previously uncharacterized roles in controlling flowering. Zm00001eb082790 is a homolog of the Arabidopsis gene *trm3*, a gene involved in plasmodesmata trafficking which exhibits a lethal loss of function phenotype and delays senescence and flowering when over expressed (Benitez-Alfonso *et al*. 2009). Our eQTL analysis suggests *trm3* is either directly regulated by or downstream of *mads1*, and a second MADS-box containing gene, *mads76* (Supplemental figure S12). The auxin response factor *arf34* (Figure 3E), which closest Arabidopsis counterpart is *Atarf6*, which, along with *Atarf8* regulates stem elongation and flower maturation (Nagpal *et al*. 2005). The maize counterpart of the Arabidopsis C2H2 transcription factor *late* (late flowering), Zm00001eb059970 was also linked to flowering time in our analysis. Increased expression of *late* in Arabidopsis results in delays in bolting and flowering consistent with the association between increased expression of the *late* homolog in maize with later flowering observed here (Figure 3F).

This study demonstrates the potential of transcriptome-wide association studies (TWAS) to accelerate the characterization and study of the genes involved in controlling variation in complex traits. Our results, with large numbers of genes relative to GWAS passing stringent false discovery rate thresholds and significant numbers of these being validated in the literature, indicate how gene expression data from large populations combined with good sequencing depth and narrow sample collection windows can generate large numbers of well supported hypotheses about the roles of individual genes in controlling individual traits. We also demonstrate the reusability of transcript abundance datasets across different environments and the ability to detect genes known to act in different tissues from the ones in which we profiled gene expression, suggesting broader potential to reuse population level expression datasets like the one described here with data on new traits scored in new environments.

## Materials and methods

### Field experiments and trait scoring

Two field studies were conducted as part of this experiment, using a common set of seed stocks. In both, large subsets of the Wisconsin Diversity Panel (Mazaheri *et al*. 2019) were grown in replicated trials conducted at the University of Nebraska-Lincoln’s Havelock Farm near Lincoln, Nebraska (40.852 N, 96.616 W) and Michigan State University’s Agronomy Farm near East Lansing, Michigan (42.709 N, 84.469 W). The experimental design and trait scoring of the Lincoln, Nebraska field trial has been previously described (Mural *et al*. 2022). Briefly, a total of 1,680 plots were grown in a randomized complete block design with each block consisting of 840 plots constituting 752 unique genotypes, plus a single repeated check genotype. In Michigan, a total of 1,520 plots were grown in a randomized complete block design with each block consisting a single plot each of 760 unique genotypes. In both environments, the date of anthesis for a given plot was considered to be the first day that at least 50% of plants in the plot had visible pollen shed. The date of silking for a given plot was considered to be the first day when visible silks were present on at least one ear shoot for at least 50% of the plants in the plot. “Days to silking” and “days to anthesis” were calculated relative to the planting dates at each location: May 6^th^ 2020 in Lincoln, Nebraska and May 25^th^, 2020 in East Lansing, Michigan (Supplemental dataset S1).

### Correcting for spatial variation in trait datasets

Corrections for spatial variation within a location was performed using the R package SpATS (Velazco *et al*. 2017) to fit a 2-dimensional penalized spline model to raw plot level trait measurements. In order to get plot-level values rather than BLUPs/BLUEs – the default output of the package – the SpATS model was fit using plot number rather than genotype name as the “genotype” input value. The model was fit using one knot per two rows and one knot per two columns. Spatially corrected plot level measurements are provided in Supplemental dataset S2.

### Repeatability analyses

A total of 751, 750, 758, and 749 genotypes with trait measurements in both replicated blocks were used to estimate repeatability for silking in Nebraska, anthesis in Nebraska, anthesis in Michigan and silking in Michigan, with the lower number of genotypes in Michigan reflecting a greater proportion of missing values in the dataset. Repeatability from anthesis and silking in Nebraska and Michigan was defined using the following formula:

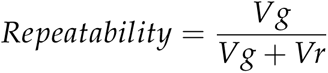

Where *V_g_*is the proportion of total variance explained by genotype and *V_r_*is the residual variance. *V_g_* and *V_r_* were estimated using the *lmer* function inside the lme4 R package to fit a simple model (spatially corrected trait = genotype effect + residual) for each trait in each environment.

Repeatability analysis for gene expression was conducted as described above, with the modification that expression data was taken from 51 genotypes where expression data was estimated twice using two separately collected biological samples from the same plots in the field.

### Quantifying Gene Expression

Tissue samples were collected on July 8^th^, 2020 from one of two replicated blocks – block 1, the western-most block – of Lincoln, Nebraska field experiment described above. Samples were collected from a single representative plant per plot, excluding edge plants where possible. Five leaf disks were collected from the pre-ante-penultimate leaf (the fourth from the highest visible and emerged leaf) of the selected plant (Figure S2A). Leaf tissue was immediately flash frozen in liquid nitrogen and then packed on dry ice until samples were loaded into a -80°C freezer. Samples were collected in parallel by seven researchers, allowing all samples to be collected over a period of approximately two hours, with all sampling completed prior to noon on the day of collection.

Frozen tissue samples were ground without a buffer suspension using a TissueLyzer II (Qiagen; 85300) that oscillated at 25Hz in 30 second increments, in a two step process, resting the samples in dry ice for one minute between grindings to ensure they were completely frozen.

RNA was extracted from the resulting ground samples using a Kingfisher Flex automated extraction robot (ThermoFisher Scientific; 5400630) and the MagMax Plant RNA Isolation Kit (ThermoFisher Scientific; A47157) following the manufacturer’s protocol. Twelve samples from each batch of 95 samples extracted in parallel were run on a 1% agarose gel and visually inspected for evidence of sample degradation to confirm the quality of extracted RNA. The RNA concentration of each sample was quantified using the Quant-IT Broad Range RNA Assay Kit (ThermoFisher Scientific; Q10213) and a CLARIOstar Plus plate reader (BMG LabTech). RNA samples were shipped to Psomagen (Rockville, MD USA) where mRNA purification, cDNA synthesis, and sequencing library construction were performed using Illumina (San Diego, CA USA) TruSeq strand-specific RNA-seq kits. Libraries were pooled and sequenced on NovaSeq 6000 Illumina Sequencers using 2x150 bp sequencing runs and a target of 20 million fragments and 6 gigabases of sequence per sample.

Raw sequence data was filtered and low-quality sequences were removed using trimmomatic (v0.33) with the following parameters: “ILLUMINACLIP: TruSeq3-PE.fa:2:30:10 LEADING:3 TRAILING:3 SLIDINGWINDOW:4:15 MINLEN:35” (Bolger *et al*. 2014). Kalliso (v0.46) was used to estimate the expression of each maize gene in each sample in units of transcripts per millions (Bray *et al*. 2016) with the “primaryTranscriptOnly” sequence file for B73_RefGen_V5 sequence file (Schnable *et al*. 2009; Hufford *et al*. 2021) provided by Phytozome (Goodstein *et al*. 2012). After removing samples with extreme values based on a principal component analysis, genes with low expression levels were filtered out.

### Linking transcript abundance to phenotype

For each gene, transcript abundance, denoted in transcripts per million, was converted to a range from 0 - 2 using the methodology described in Li *et al*. (2021). Briefly, to minimize the effect of extreme values in individual samples, the 5% of samples with the lowest transcripts per million values for each gene were scored as 0, the the 5% of samples with the highest transcripts per million values for each gene were scored as 2, and the remaining 90% of samples were re-scaled between 0 and 2 using the formula:

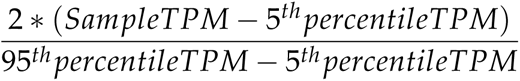

TWAS was performed using the compressed mixed linear model as implemented in GAPIT (v3.1) to link gene expression, normalized as described above, with variation in spatially normalized measurements of male and female flowering time in Nebraska and Michigan (Zhang *et al*. 2010; Lipka *et al*. 2012). The three first principal components of variation calculated by GAPIT from the expression data were included as covariates. The threshold for a statistically significant association between transcripts and phenotypic variation was calculated independently for each trait at a p-value corresponding to a false discovery rate of 0.05 calculated using the Benjamini & Hochberg method (Benjamini and Yekutieli 2001).

### Linking genetic markers to phenotype

Genome-wide association was conducted using a published resequencing-based genetic marker data for 752 genotypes drawn from the Wisconsin Diversity panel (Grzybowski *et al*. 2023). This marker set had already been filtered to exclude markers with unusually high or low sequencing depth indicative of copy number variants, as well as for markers with *≥* 50% missing data and had been imputed using Beagle 5.0 (Browning *et al*. 2018). The dataset was further filtered to retain only markers with minor allele frequency >0.05 among the 693 genotypes included in this study using plink2 (v2.0a1) (Chang *et al*. 2015), resulting in a final dataset of 15,659,765 genetic markers. Significant associations between filtered genetic markers and the same phenotype values employed for TWAS were identified using the linear mixed model as implemented in GEMMA (v0.98) (Zhou and Stephens 2012) with three principal components of variation and a kinship matrix – previously calculated from the genetic markers using plink2 and GEMMA, respectively (Chang *et al*. 2015; Zhou and Stephens 2012) – included as covariates.

### Linking genetic markers to gene expression of the candidate genes

eQTL mapping was performed using the mixed linear method implemented within rMVP (V1.0.6) (Price *et al*. 2006; Yin *et al*. 2021) Box-Cox transformed TPM estimates of candidate gene expression across the population of 693 individuals previously transformed with the Box-Cox method (Osborne 2010) and the same set of genetic markers described above. Three principal components of variation and a kinship matrix calculated using the VanRaden method (VanRaden 2008) were included as covariates. Linkage disequilibrium analysis was conducted using plink 1.9 (Purcell *et al*. 2007).

## Data availability

RNA-Seq data for all lines used in this study is available from the European Nucleotide Archive (ENA) under the study accession number: PRJEB67964. Big gene expression calculated as transcript per million is public in the GitHub repository: https://doi.org/10.6084/m9.figshare.24470758.v1

## Acknowledgments

The authors would like to thank: Alice Guo, Elijah Frost, Isaac Stevens, Leighton Wheeler, Nathaniel Pester, Thomas Hoban and Christine Smith for their work planting and maintaining the Nebraska field and collecting trait data and tissue samples. The authors would also like to thank: Sidney Sitar, Sophia Holtz, Carolina Freitas, Emma Chrzanowski, Matthew Salo, Lauren Truitt, Eli Hugghis, Blake Trygestad, Robert Shrote, and Zhongjie Ji for their work maintaining the Michigan field site and collecting trait data, as well as Ruijuan Tan for her help in assembling and curating the Michigan trait dataset. This work was conducted in part using the Holland Computing Center of the University of Nebraska, which receives support from the Nebraska Research Initiative.

## Author Contributions

JVT-R and JCS conceived of the project. AMT and LN generated the common seed stocks used for both trials. JT, LN, LL-C, AMT and JCS generated, assembled, and quality controlled data. DL, JT, JD, GS, MWG and RVM designed and advised on analysis methods. JVT-R, JD, WA, RVM conducted analyses. JVT-R, LL-C, and WA visualized the results. JVT-R, RVM and JCS wrote the first draft of the manuscript. DL, JT, LN, LL-C, WA, MWG and AMT contributed significant additional content during the revision of the manuscript. All authors read and approved the final version of the manuscript.

---

**Figure S1.**
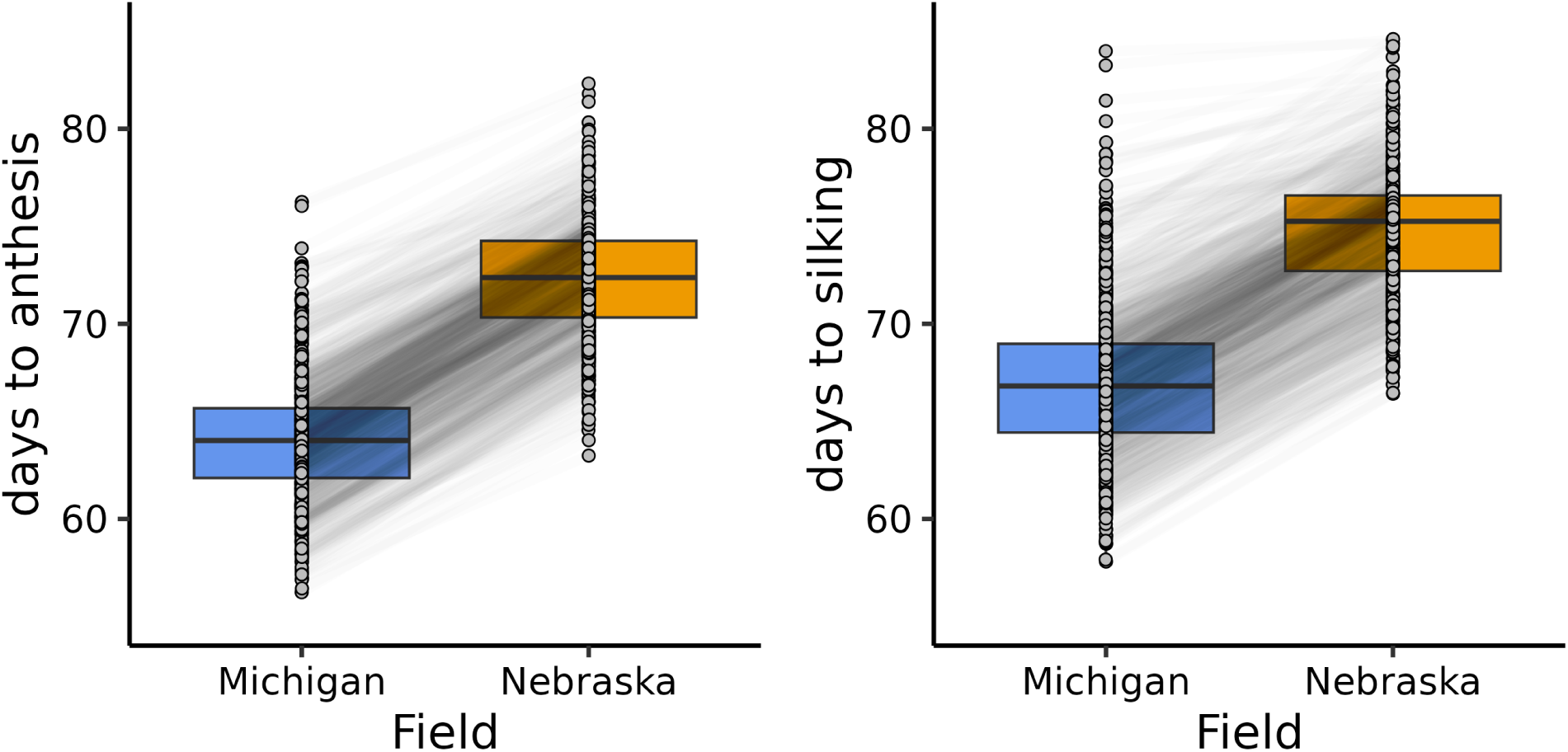
Flowering time from 699 lines used for TWAS and GWAS in this study. Left panel shows the distribution of days to anthesis in Michigan and Nebraska. Right panel shows the distribution of days to silking in Michigan and Nebraska. single genotypes are represented with grey dots and their respective genotype in the other environment is linked with a straight line.

**Figure S2.**
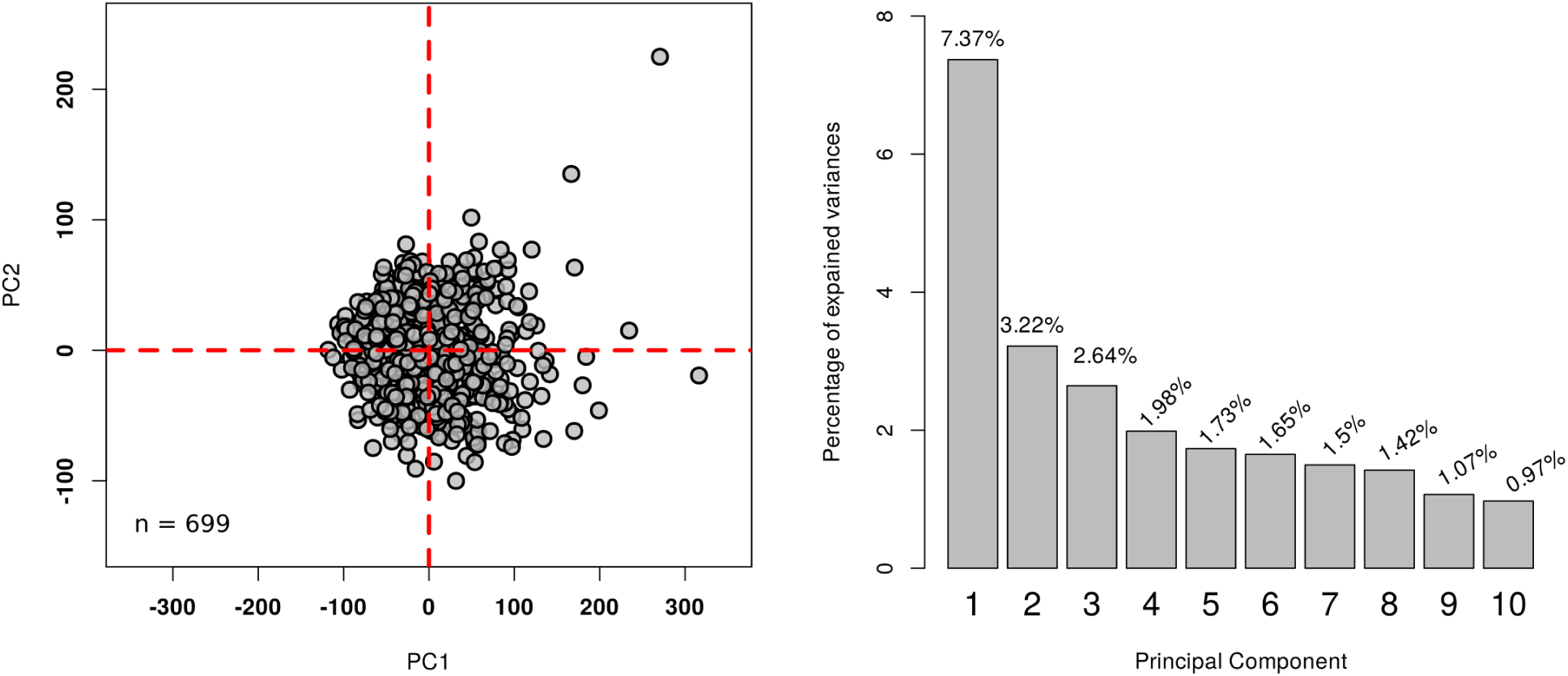
Principal Component Analysis of 699 lines. Left panel shows the distribution of 699 genotypes based on the expression reported as transcripts per million (TPM) of 39,756 genes before removing low expressed genes. Right panel showed the percentage of variance explained by the first 10 principal analysis.

**Figure S3.**
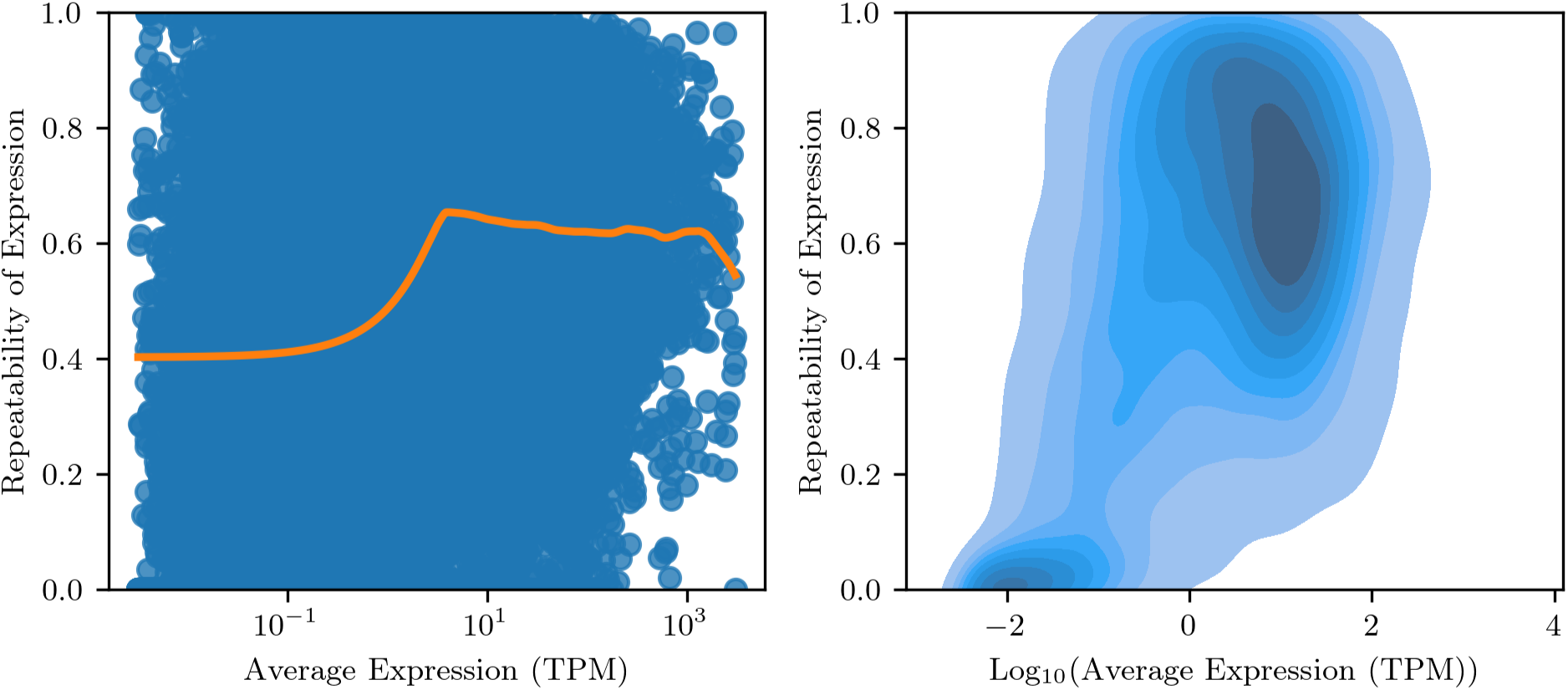
Repeatability of genes according to their expression. Left panel shows the distribution the average expression in TPM of 39,756 genes calculated from two replicates and their respective repeatability. Right panel shows distribution of the log10 transformed average expression in TPM of 39,756 genes calculated from two replicates and their respective repeatability. Intensity of the colors indicates the density of the data points.

**Figure S4.**
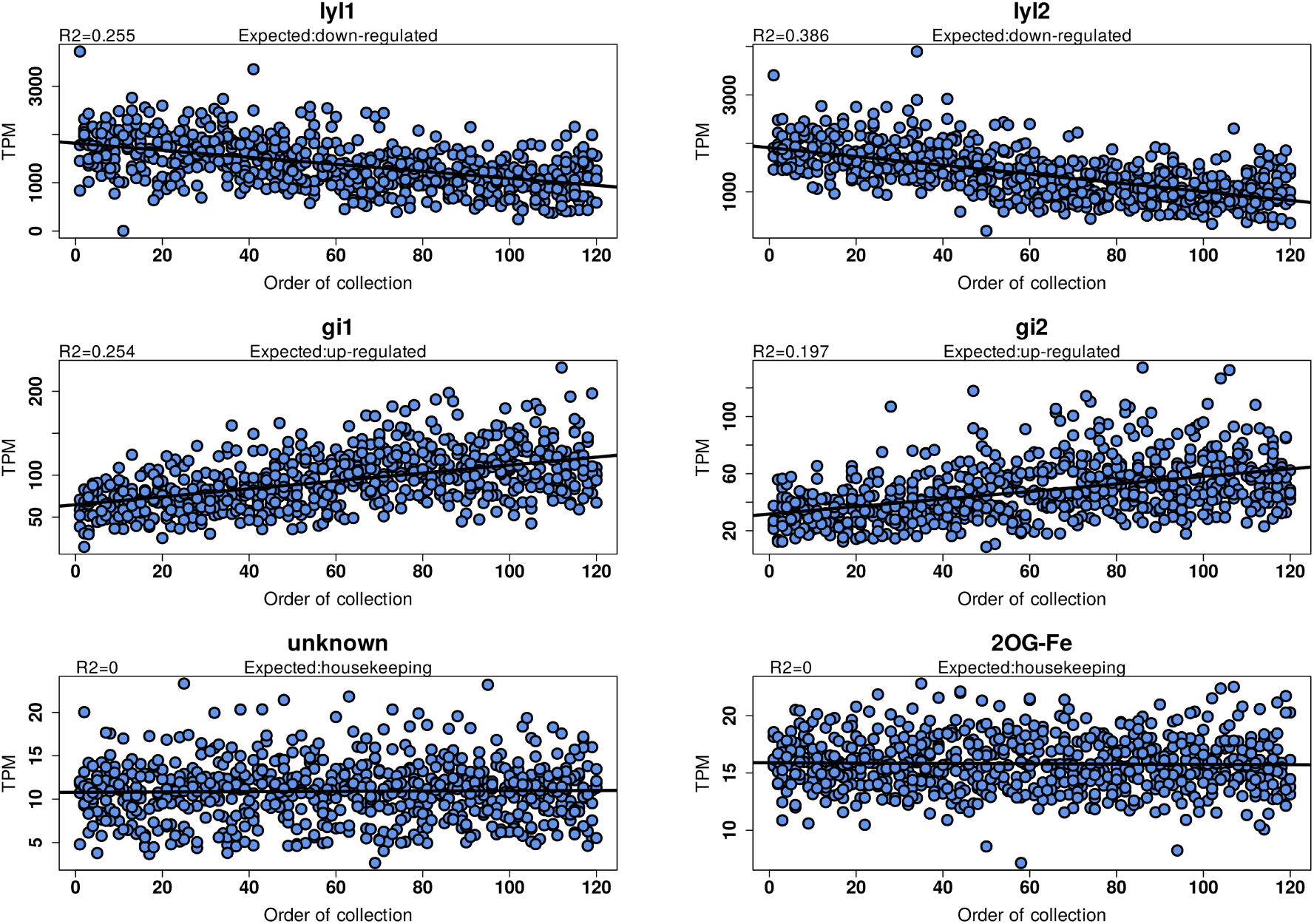
Pattern of diurnal genes. Top panel shows *lhy-like 1* (*lyl1*) and *lhy-like 2* (*lyl2*), two genes with expected down-regulation during day time. Middle panel shows *gigantea 1* (*gi1*), and *gigantea 2* (*gi2*) with expected up-regulation during day time. Bottom panel shows two housekeeping genes, *unknown* (Zm00001eb270840) and *2OG-Fe* (Zm00001eb377750). R^2^ values are showed in the corner of each plot.

**Figure S5.**
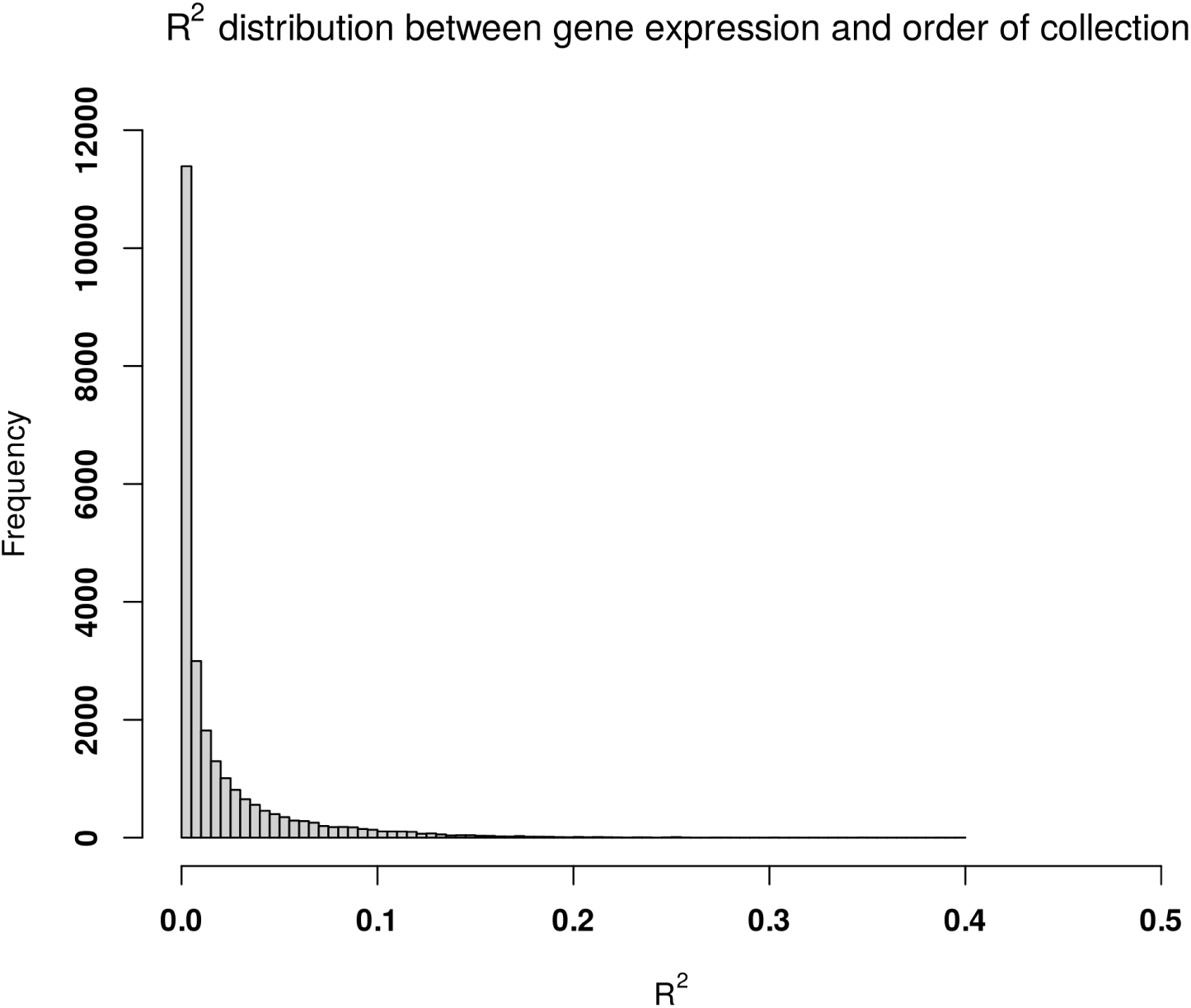
Distribution of R^2^ between gene expression and the order of collection. R^2^ for the expression of 24,585 genes used in this study and the order of collection.

**Figure S6.**
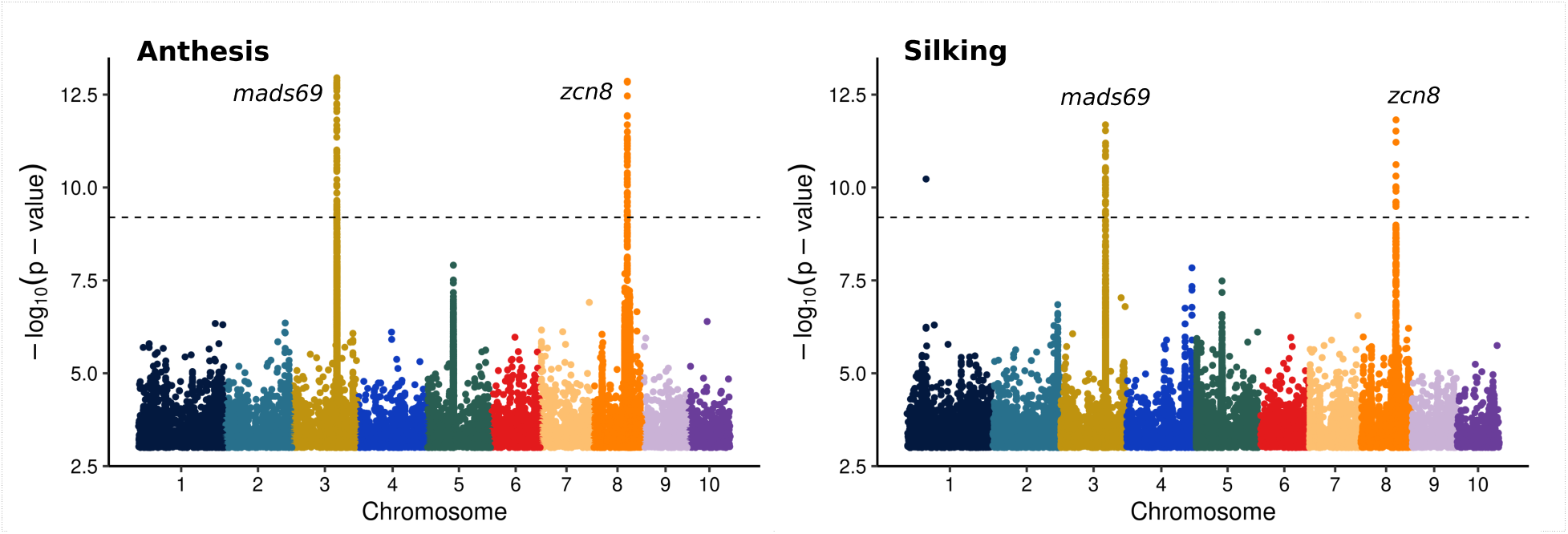
Genes associated with days to anthesis and days to silking in Nebraska via GWAS. Left panel shows a Manhattan plot of days to anthesis using GWAS. The horizontal line represents a Bonferroni correction cutoff of 0.05, which assumes all markers as independent tests, n = 15,659,765.

**Figure S7.**
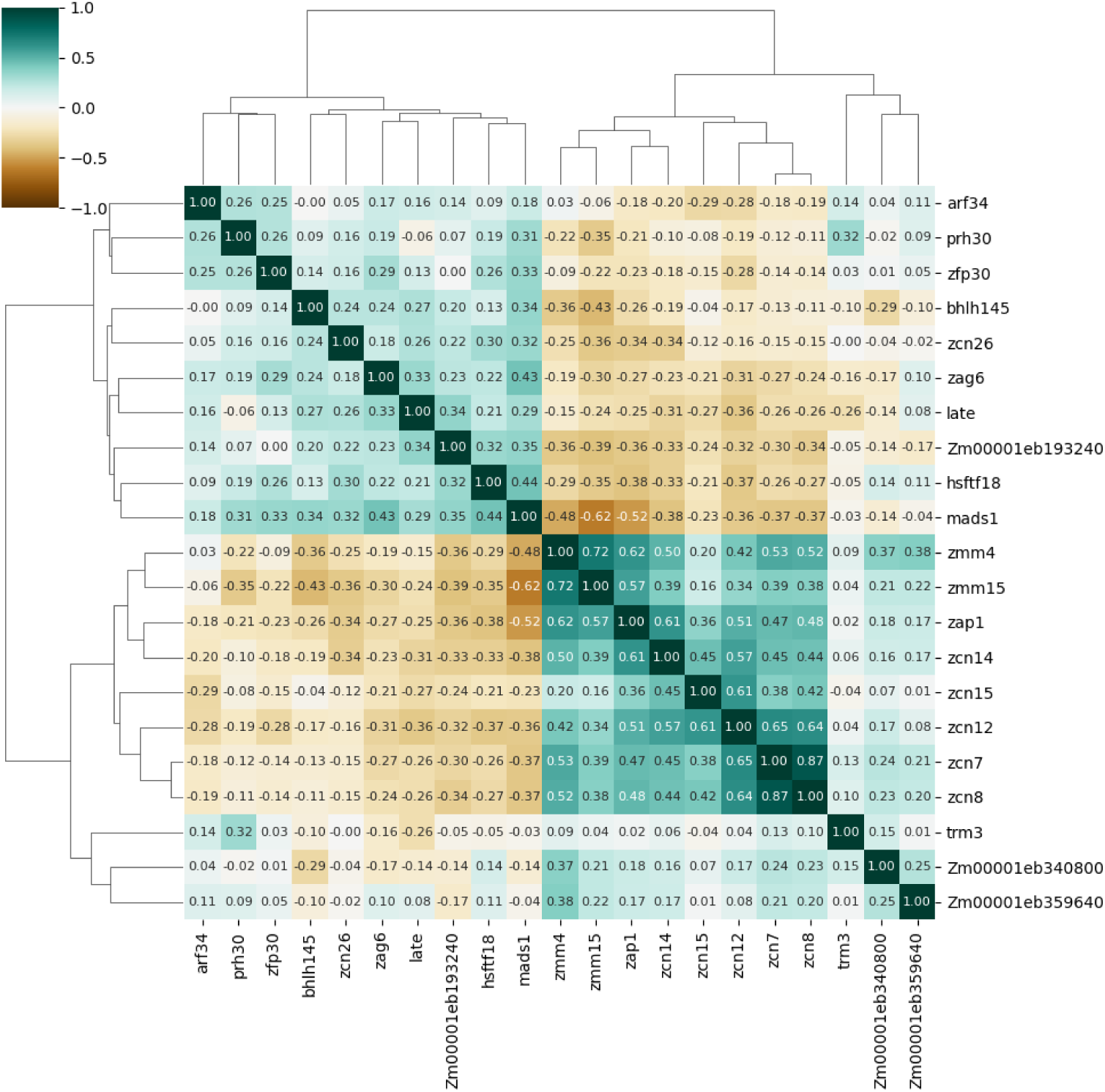
Spearman correlation of the expression of associated genes via TWAS. Positive correlation is colored in green while negative correlation is shown in red.

**Figure S8.**
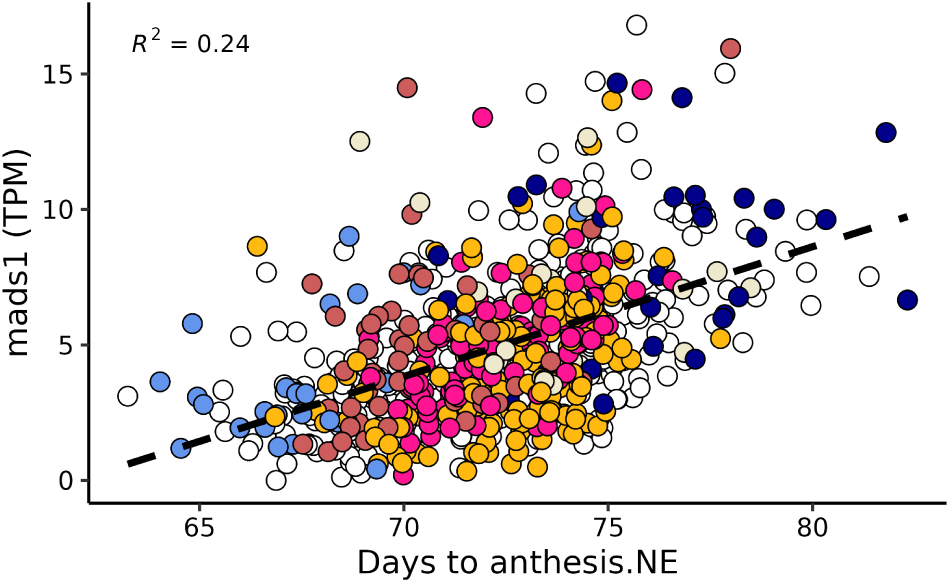
Correlation of *mads1* with flowering time in Nebraska. Left panel shows the correlation of transcripts per million (TPM) of *mads1* with days to anthesis represented from Nebraska field.

**Figure S9.**
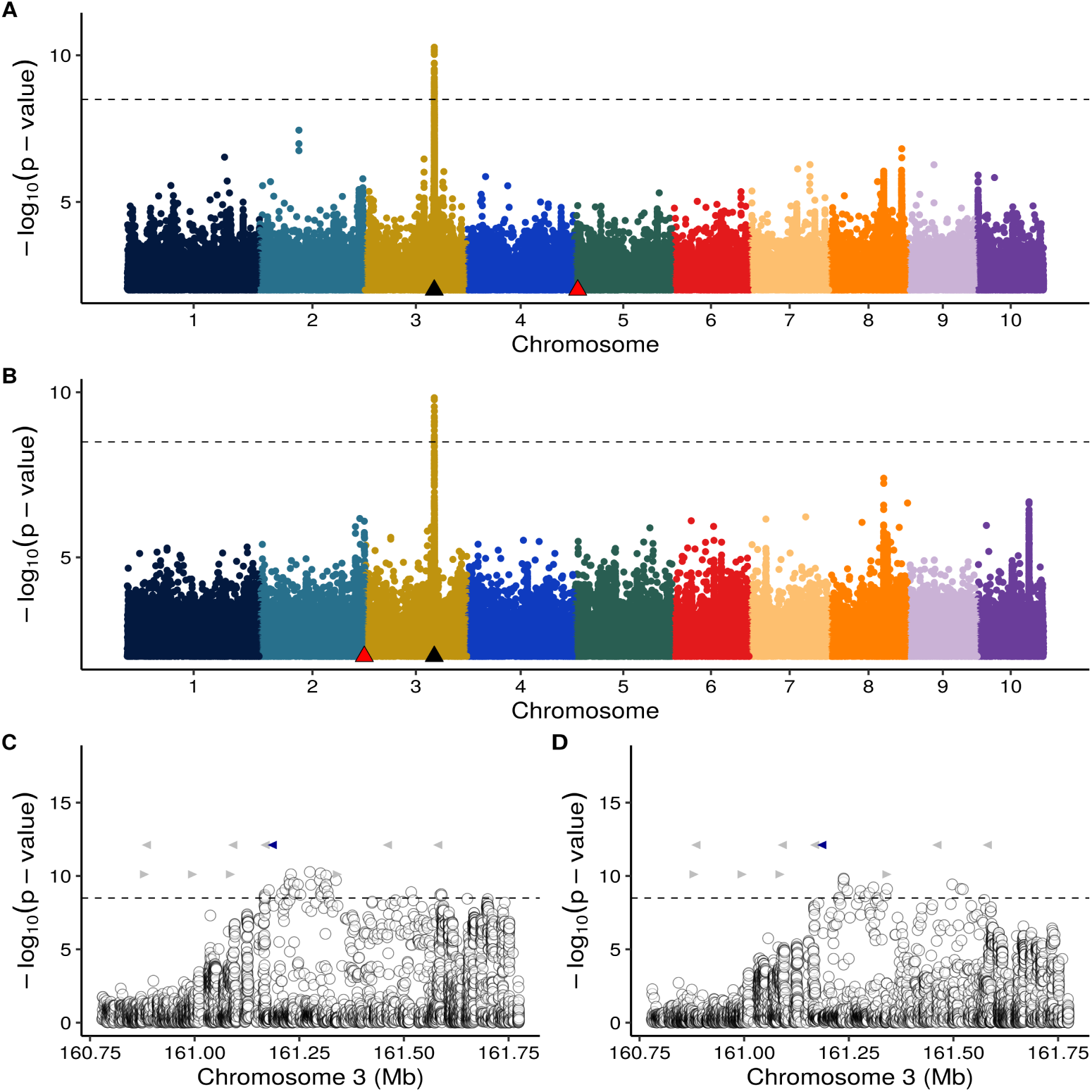
*Trans* eQTL upstream *mads69* are associated with the expression of *zmm15* and *zap1*. Genome-wide association analysis of the expression of **A** *zmm15* and **B** *zap1*. Analysis was conducted using 15,659,765 genetic variants. Red triangle at the bottom of the dots in chromosome 5 and chromosome 2 represent the position of *zmm15* and *zap1*, respectively. Black triangle at the bottom of the dots in chromosome 3 represents *mads69*. Horizontal dashed line indicates a 0.05 threshold after Bonferroni correction which assumes all variants are independent tests. **C** Zoomed in view of the peak located on chromosome 3. Blue triangle indicates the position of *mads69*. Data from eQTL analysis of *zmm15*. **C** Zoomed in view of the peak located on chromosome 3. Blue triangle indicates the position of *mads69*. Data from eQTL analysis of *zap1*

**Figure S10.**
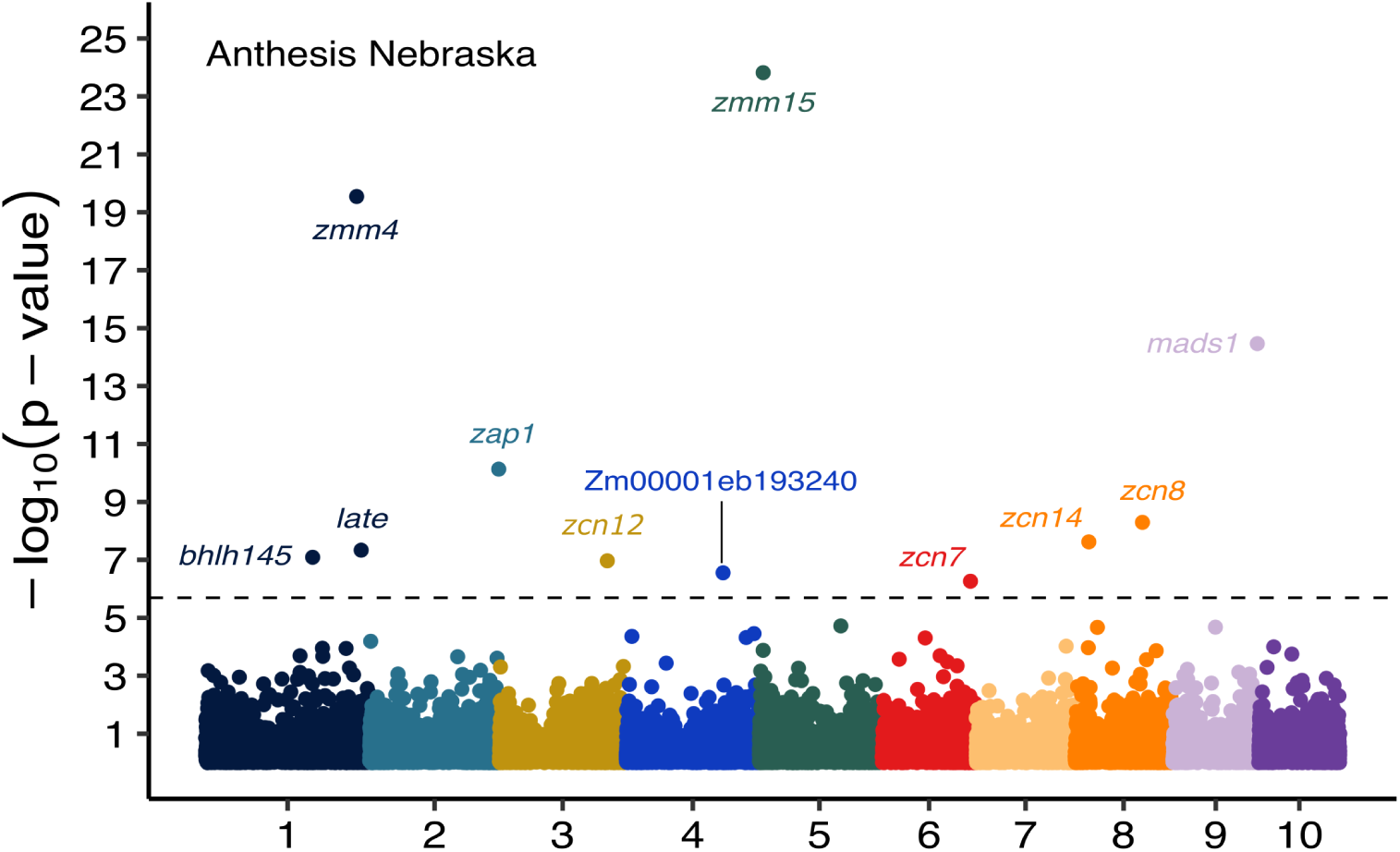
Bonferroni correction applied to days to anthesis recorded in Nebraska. The dashed line represents the threshold after Bonferroni correction of 0.05 assuming each gene as an independent test, n = 24,585.

**Figure S11.**
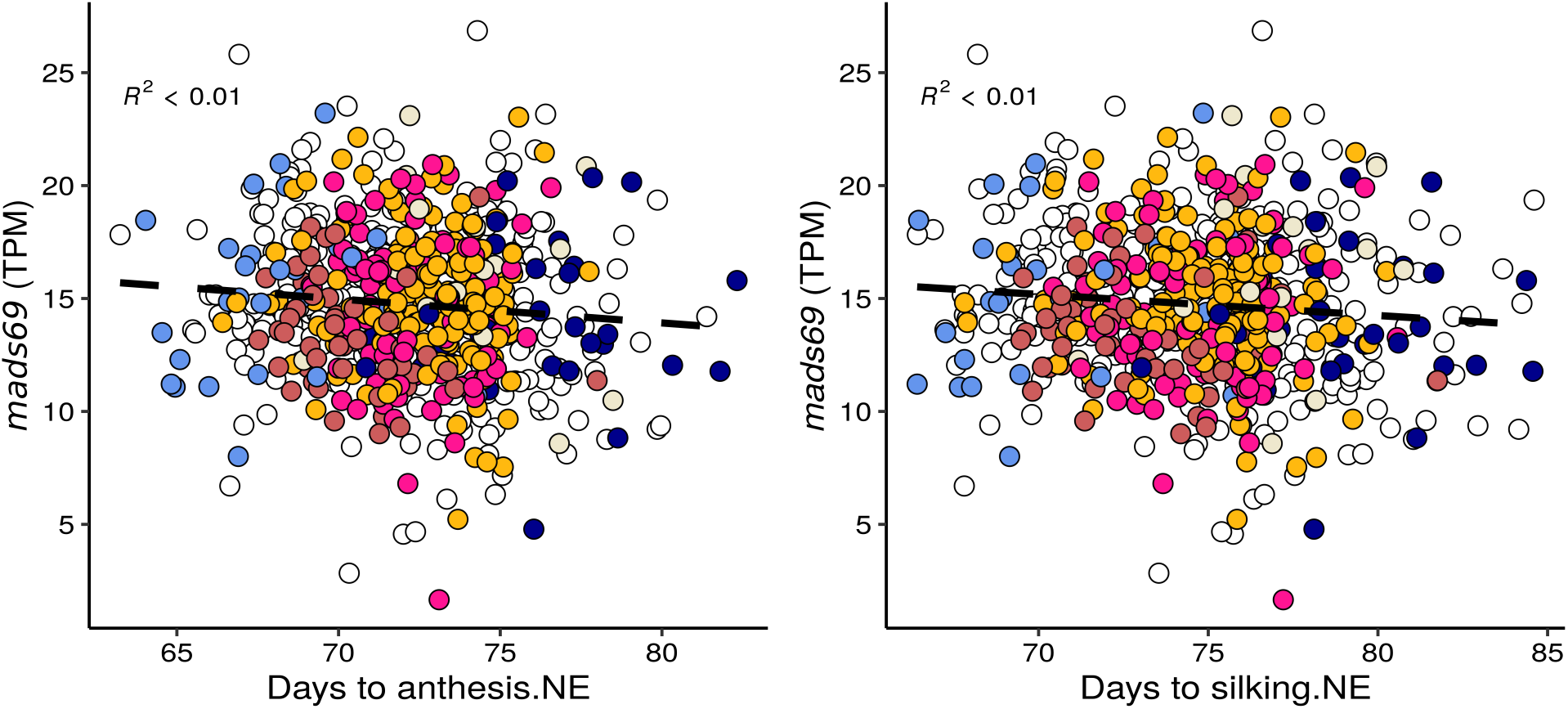
Correlation of *mads69* with flowering time in Nebraska. Left panel shows the correlation of transcripts per million (TPM) of *mads69* with days to anthesis measured from Nebraska field. Left panel shows the correlation of transcripts per million (TPM) of *mads69* with days to silking from Nebraska field.

**Figure S12.**
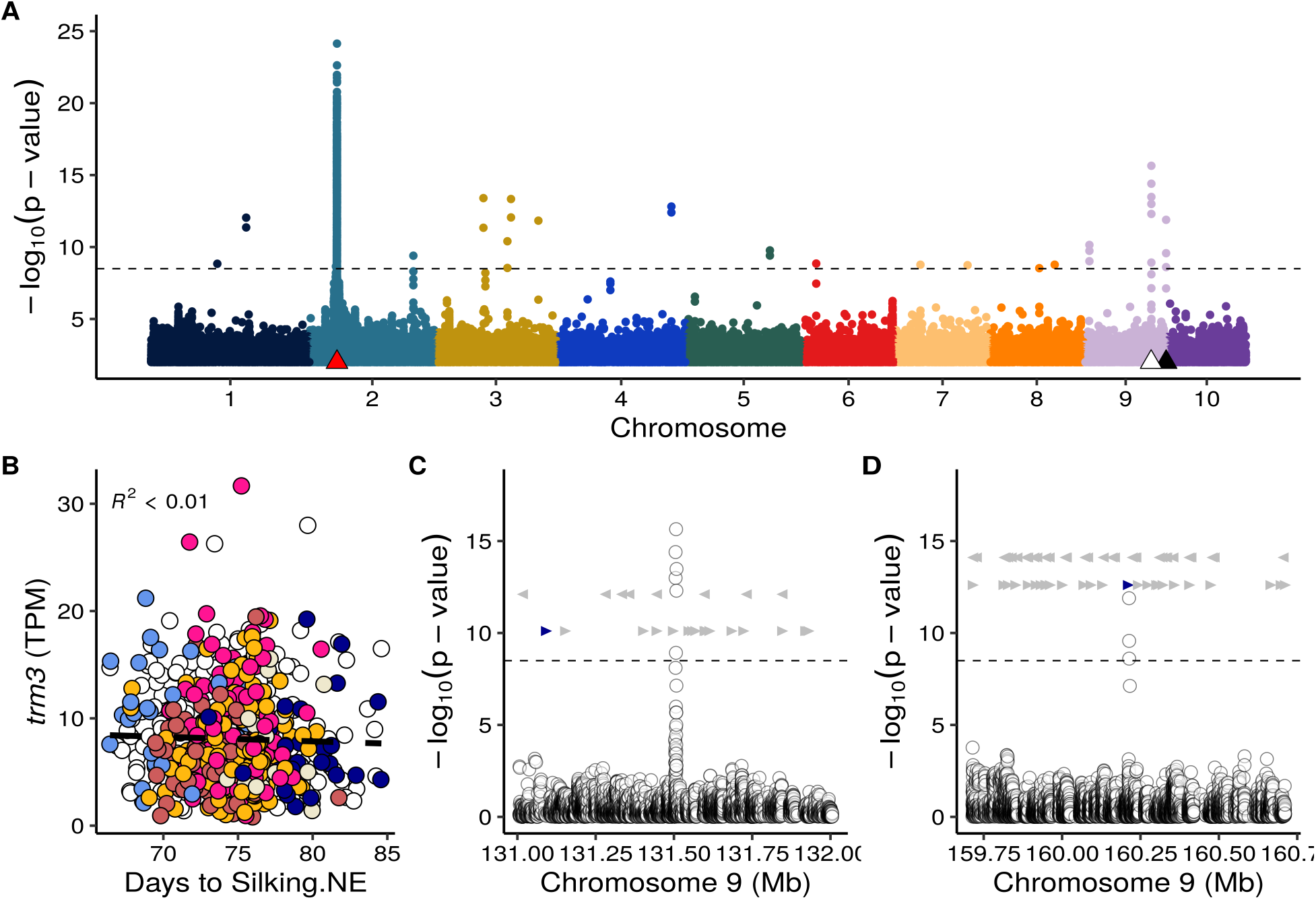
Flowering time genes in *trans* are associated with *trm3* expression. **A** Genome-wide association analysis of the expression of *trm3*. Analysis was conducted using 15,659,765 genetic variants; for further information see Materials and Methods section. Red triangle at the bottom of the dots in chromosome 2 represents the position of *trm3*. White triangle at the bottom of the dots in chromosome 9, represents *mads76*. Black triangle at the bottom of the dots in chromosome 9, represents *mads1*. Horizontal dashed line indicates a 0.05 threshold after Bonferroni correction which assumes all variants are independent tests. **B** Relation of *trm3* expression reported as TPM with days to silking scored from Nebraska 2020. Dots are colored based on sub-groups referred to in Figure 1. The Linear dashed line indicates the calculated regression line. **C** Zoomed in view of the peak located on chromosome 9. Blue triangle indicates the position of *mads76*. **C** Zoomed in view of the peak located on chromosome 9. Blue triangle indicates the position of *mads1*.

**Table S1** Flowering time data scored from Nebraska 2020 and Michigan 2020.

**Table S2** Spatial corrected and metadata for the 693 lines used in this study.

**Table S3** Correlation between PCs and sampling time.

**Table S4** Origin of lines based on the country of origin, modified from (Grzybowski *et al*. 2023).

